# Delivery of caspase inhibitors through GSDMD pores to inhibit pyroptosis

**DOI:** 10.1101/2025.02.11.637513

**Authors:** Katarzyna M. Groborz, Melissa E. Truong, Irma Stowe, Bettina Lee, Robert S. Jones, Emile Plise, Elizabeth S. Levy, Ponien Kou, Wyne P. Lee, Juan Zhang, Julia Nguyen, Malgorzata Kalinka, Marcin Drag, Nobuhiko Kayagaki, Kim Newton, Marcin Poreba, Vishva M. Dixit

## Abstract

Caspase-1, -4, -5, and -11 activate Gasdermin D (GSDMD) to form pores in the plasma membrane. In addition to releasing interleukin (IL)-1β and IL-18, GSDMD pores cause a lytic, proinflammatory form of cell death called pyroptosis. Blocking this pathway holds therapeutic promise for the treatment of inflammatory disorders, but clinical trials of cell permeable caspase inhibitors have been unsuccessful. Here, we describe covalent inhibitors of proinflammatory caspases that are impermeable to healthy cells but effectively block caspase-1 driven pyroptosis and IL-1β secretion. Their failure to inhibit apoptosis implies inhibitor entry through GSDMD pores. Propidium iodide entered rescued cells, confirming transient membrane permeabilization via caspase-1 and GSDMD. Caspase-1 inhibition prevented rather than delayed cell death, likely due to the activation of membrane repair mechanisms neutralizing the initial GSDMD pores. Notably, inhibiting caspase-1 and -11 suppressed IL-1β and IL-18 production in a mouse model of endotoxic shock. These findings underscore the therapeutic potential of exploiting GSDMD pores for the delivery of caspase inhibitors, offering a novel strategy for treating inflammatory diseases.

---

Cell death in multicellular organisms plays pivotal roles in development, homeostasis, and pathogen defense ^1^. The programmed cell death pathways of apoptosis and pyroptosis are mediated by aspartate-specific cysteine proteases called caspases. The so called proinflammatory caspases (human caspase-1, -4, and -5; mouse caspase-1 and -11) cleave and thereby activate the pore-forming protein gasdermin D (GSDMD) to elicit pyroptosis, a lytic form of cell death. Distinct caspases, including caspase-8 and -9, initiate apoptosis by cleaving and activating caspase-3 and - 7. In excess, both apoptosis and pyroptosis have been shown to cause inflammatory disorders in mice ^1^. The potential therapeutic benefit of caspase inhibition in human diseases that are scharacterized by uncontrolled cell death remains uncertain. Clinical trials of cell permeable pan-caspase inhibitors have failed owing to toxicity or a lack of efficacy ^2^. One potential liability of pan-caspase inhibition is that inactivation of caspase-8 sensitizes certain cell types to a lytic form of cell death termed necroptosis ^1^. The pursuit of caspase-selective inhibitors has been challenging because of pronounced structural homologies among the caspases ^3^. Recently, extracellular nanobodies against GSDMD or the caspase-1 adaptor ASC were shown to prevent pyroptosis, presumably entering cells through the initial GSDMD pores and arresting further pore assembly ^4,5^. A pause in GSDMD pore formation was posited to give the membrane repair machinery time to resolve damage to the cell membrane ^4^. These findings prompted us to test whether typically membrane-impermeable caspase inhibitors with selectivity for caspase-1, -4, -5, and -11 might also enter cells through GSDMD pores and prevent pyroptosis. If these caspase inhibitors only enter cells that have engaged the pyroptosis pathway then they might be tolerated better than cell permeable pan-caspase inhibitors.

## Substrate specificity of caspase-1/4/5

As a first step towards generating libraries of potentially selective caspase inhibitors, we screened the substrate preferences of human caspase-1, -4, and -5 using hybrid combinatorial substrate libraries (HyCoSuL) incorporating diverse tetrapeptides with Asp in the P1 position (Extended Data Fig 1). HyCoSuL sublibraries P2, P3 and P4 were composed of tetrapeptides with an equimolar mixture of natural amino acids in two positions (e.g. P2 and P3) and a fixed natural or unnatural amino acid in the remaining space (e.g. P4) ^6^. Human caspase-1, -4, and -5 resembled mouse caspase-1 and -11 ^7^ by favouring His in the P2 position (Extended Data Fig. 1). By contrast, human caspase-3 prefers small, aliphatic P2 residues like Val and Ile ^8^. All five caspases preferred Glu in the P3 position, but caspase-11 better tolerated bulky residues like Bip, Bpa, and Tyr (2,6-Cl2-Bzl) (Extended Data Fig. 1) ^6^. Caspase-11 also accommodated small, aliphatic P4 residues (Tle, Val and Ile), whereas caspase-1, -4, and -5 favoured bulky P4 residues (Trp and Tyr). Thus, human caspase-4 and mouse caspase-11 have distinct cleavage site preferences, despite being the core component of the non-canonical inflammasome that responds to cytoplasmic lipopolysaccharide (LPS) in both species ^9,10^.

Given that human caspase-1, -4, and -5 have similar cleavage site preferences, we tested their ability to cleave a set of tetrapeptide caspase substrates described previously ^7^. A tetrapeptide bearing the Trp-Val-His-Asp sequence found in human pro-IL-1β was among those most susceptible to cleavage by human caspase-1, whereas hydrolysis of the Phe-Leu-Thr-Asp sequence found in human GSDMD was approximately 3-fold less efficient (Extended Data Table 1). Interestingly, caspase-1 exhibited catalytic efficiencies (k_cat_/K_M_) that were an order of magnitude greater than those of caspase-4, -5, and -11 (Extended Data Table 1) ^7^. The incorporation of non-natural amino acids into the tetrapeptide sequences elevated the cleavage efficiencies, while concurrently enhancing the selectivity factor for proinflammatory over proapoptotic caspases.

## Screening a caspase inhibitor library

Taking into account the cleavage site preferences of caspase-1, -4, -5, and -11, we synthesized around 100 tetrapeptides bearing an acyloxymethyl ketone (AOMK) reactive group to covalently modify their targets in an irreversible manner ^11,12^. We determined their inhibition parameters on recombinant human caspase-1, -3, -4, -6, -7, -8, -9, and -10 (Extended Data Table 2). Tetrapeptides KGR-3, KGR-23, and KGR-72 stood out as potent and somewhat selective inhibitors of human caspase-1 (Extended Data Table 2), and were subsequently found to also inhibit mouse caspase-1 (Extended Data Table 3). The cell permeability of KGR-3, KGR-23, and KGR-72 was deemed very low when assessed in gMDCKI cells lacking the efflux pump MDR1 ^13^ (Extended Data Table 4).

To determine if any of our tetrapeptides might pass through GSDMD pores in the cell membrane to inhibit caspase-1, we tested their ability to block NLRP3- and caspase-1-dependent pyroptosis in LPS-primed human THP-1 monocytic cells treated with nigericin. Cell lysis was monitored by lactate dehydrogenase (LDH) release. As expected, nigericin-induced cell lysis was inhibited by the NLRP3 inhibitor MCC950 ^14–16^, the pan-caspase inhibitor emricasan (IDN-6556) ^17^, caspase-1 deficiency, or GSDMD loss (Extended Data Fig. 2a). Several tetrapeptides, including KGR-3, KGR-23, and KGR-72, greatly reduced or inhibited nigericin-induced cell lysis (Extended Data Fig. 2b).

We also screened a subset of our tetrapeptides for their ability to block caspase-4- or -11-driven pyroptosis, focusing on those with the highest potency against recombinant caspase-1 or that best blocked NLRP3-dependent pyroptosis in THP-1 cells. Rather than electroporate or transfect LPS into cells to activate caspase-4 or -11 ^10,18,19^, we engineered THP-1 cells to express either FK506 binding protein 12 (FKBP)-caspase-4 or FKBP-caspase-11 fusions where FKBP replaced the caspase pro-domain. Cell permeable B/B ligand was added to dimerize the FKBP fusions, resulting in GSDMD cleavage and pyroptosis that was inhibited by the pan-caspase inhibitor emricasan (Extended Data Fig. 3a-d). We identified several tetrapeptides that were comparable to emricasan in suppressing cell death induced by caspase-4 and/or -11 (Extended Data Fig. 3d). For example, KGR-3 blocked GSDMD processing and cell death induced by caspase-4, but was less effective against caspase-11 (Extended Data Fig. 3d). Others, including KGR-23, KGR-53, and KGR-72, suppressed cell death induced by caspase-4 or -11. To exclude tetrapeptides that might access cells in a GSDMD-independent manner and be less selective for caspase-1, -4 or -11, we also screened for inhibition of staurosporine-induced apoptosis, which is thought to be dependent on BAX or BAK in the mitochondrial pathway that activates caspase-9 ^20^. However, unlike emricasan, none of the selected tetrapeptides suppressed staurosporine-induced cell death (Extended Data Fig. 3e).

## Characterization of inhibitor KGR-3

It is difficult to distinguish apoptotic and pyroptotic morphologies in THP-1 cells owing to their small size, so we also used human endothelial-like EA.hy926 cells to characterize KGR-3, one of the most potent caspase-1 inhibitors. EA.hy926 cells treated with Val-boroPro (VBP) to trigger the NLRP1 or CARD8 inflammasomes ^21–23^ exhibited caspase-1- and GSDMD-dependent swelling that resulted in cell lysis and nuclear staining by the membrane-impermeable dye propidium iodide (PI) (Fig. 1a; Extended Data Fig. 4). Some *GSDMD* knockout (KO) cells still appeared to die in response to VBP (Fig. 1a), reminiscent of caspase-1-driven apoptosis in *Gsdmd* KO mouse macrophages ^24,25^. Indeed, we detected apoptotic blebbing in VBP-treated *GSDMD* KO, but not *CASP1* KO EA.hy926 cells (Extended Data Fig. 4). Blebbing was followed by secondary membrane damage as measured by PI uptake. Therefore, caspase-1 can trigger pyroptosis or apoptosis in EA.hy926 cells, but pyroptosis may be more rapid than apoptosis.

**Figure 1.**
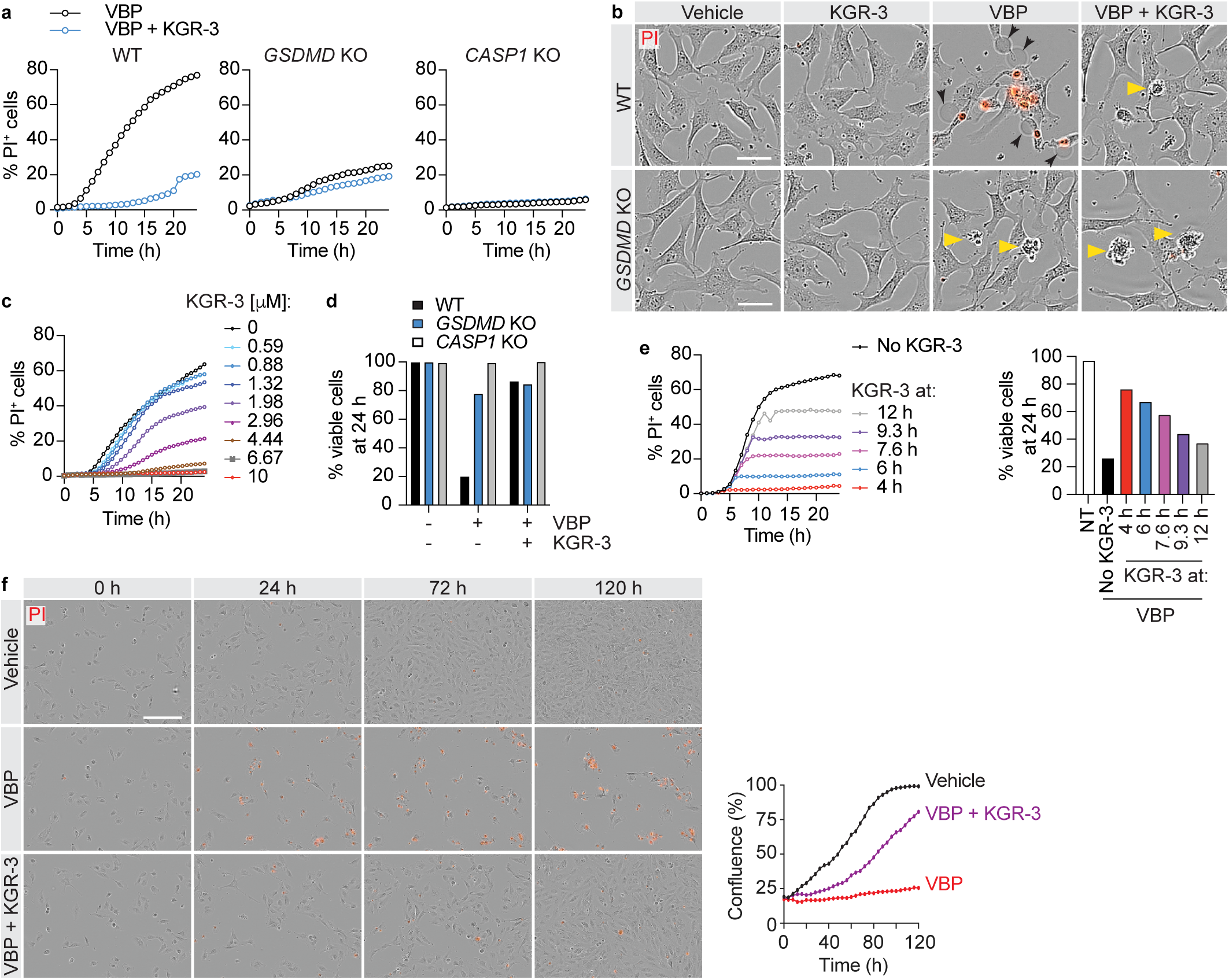
KGR3 inhibits caspase-1-dependent pyroptosis in EA.hy926 cells. **a,** Percentage of EA.hy926 cells exhibiting nuclear PI staining after treatment with 20 μM VBP ± 10 μM KGR-3. Results representative of 3 independent experiments. **b,** EA.hy926 cells treated with 20 μM VBP ± 10 μM KGR-3 in the presence of PI for 15 h. Scale bar, 40 μm. Yellow arrows highlight blebbing apoptotic cells. Black arrows indicate ballooning pyroptotic cells. PI^+^ cells are red. Results representative of 3 independent experiments. **c,** Percentage of PI^+^ WT EA.hy926 cells after treatment with 20 μM VBP. Results representative of 3 independent experiments. **d,** Percentage of viable WT EA.hy926 cells by CellTiter-Glo assay after treatment with 20 μM VBP ± 10 μM KGR-3. Results representative of 3 independent experiments. **e,** Percentage of WT EA.hy926 cells exhibiting nuclear PI staining after treatment with 20 μM VBP (left), where 10 μM KGR-3 was added at the times indicated. The percentage of viable cells was determined at 24 h (right). Results representative of 3 independent experiments. **f,** WT EA.hy926 cells treated with 20 μM VBP ± 10 μM KGR-3. PI^+^ cells are red. Scale bar, 200 μm. Results representative of 2 independent experiments.

KGR-3 suppressed VBP-induced caspase-1- and GSDMD-dependent pyroptosis in WT EA.hy926 cells (Fig. 1a, b), exhibiting a half-maximal inhibitory concentration (IC_50_) in cells of approximately 2.5 μM (Fig. 1c). A CellTiter-Glo assay, which measures cellular ATP content, confirmed that KGR-3 preserved cell viability (Fig. 1d). Importantly, however, caspase-1-dependent apoptotic blebbing in VBP-treated WT and *GSDMD* KO EA.hy926 cells was immune to KGR-3 (Fig. 1b). The ability of KGR-3 to prevent caspase-1-dependent pyroptosis, but not caspase-1-dependent apoptosis, strongly suggests that KGR-3, like other tetrapeptide caspase inhibitors with poor membrane permeability ^26^, requires GSDMD pores to access caspase-1 in the cytoplasm. Available evidence suggests that pyroptosis occurs when the assembly of GSDMD pores outpaces their elimination from membranes by the ESCRT machinery ^4,27^. Inhibition of caspase-1 by KGR-3 must favour membrane repair such that cell death is averted. By contrast, apoptotic cells that do not form GSDMD pores (for reasons that are unclear in WT cells) exclude KGR-3 and are not protected. Remarkably, addition of KGR-3 to WT EA.hy926 cells several hours after VBP treatment arrested further cell lysis (Fig. 1e). Moreover, cell death was prevented rather than just delayed by KGR-3 because the cells continued to grow over the course of 5 days, whereas few cells emerged from the control cultures exposed to VBP alone (Fig. 1f). Thus, the asynchronous nature of lethal GSDMD pore formation and pyroptosis provides a window for intervention after the initial death stimulus.

KGR-3 was also able to suppress GSDMD cleavage as well as GSDMD-dependent cell death, swelling, and lysis in B/B-treated cells expressing FKBP-caspase-4 (Extended Data Fig. 5a-d). Cell death induced by FKBP-caspase-4 was more rapid than that induced by VBP, with most of the cells dying within 6 h (Extended Data Fig. 5d). Consistent with it having low permeability in apoptotic cells, KGR-3 did not affect Tumor Necrosis Factor Related Apoptosis-Inducing Ligand (TRAIL)-induced caspase-8-dependent apoptosis in EA.hy926 cells (Extended Data Fig. 5e, f).

## Small molecule uptake via GSDMD pores

To buttress the notion that inhibitors like KGR-3 enter living cells through GSDMD pores, we determined whether small amounts of membrane impermeable, fluorescent small molecules like PI enter the cytoplasm of live cells in a GSDMD-dependent manner. When cultured in 10 μg/ml PI (which is 10 times more PI than routinely used to monitor plasma membrane integrity), EA.hy926 cells treated with VBP and KGR-3 had detectable PI in the cytoplasm (Fig. 2). Importantly, VBP-induced PI uptake in the presence of KGR-3 required both caspase-1 and GSDMD. These data suggest that the initial GSDMD pores compromise the integrity of the cell membrane towards small molecules like PI, but inhibition of caspase-1 prevents the irrevocable commitment to pyroptosis.

**Figure 2.**
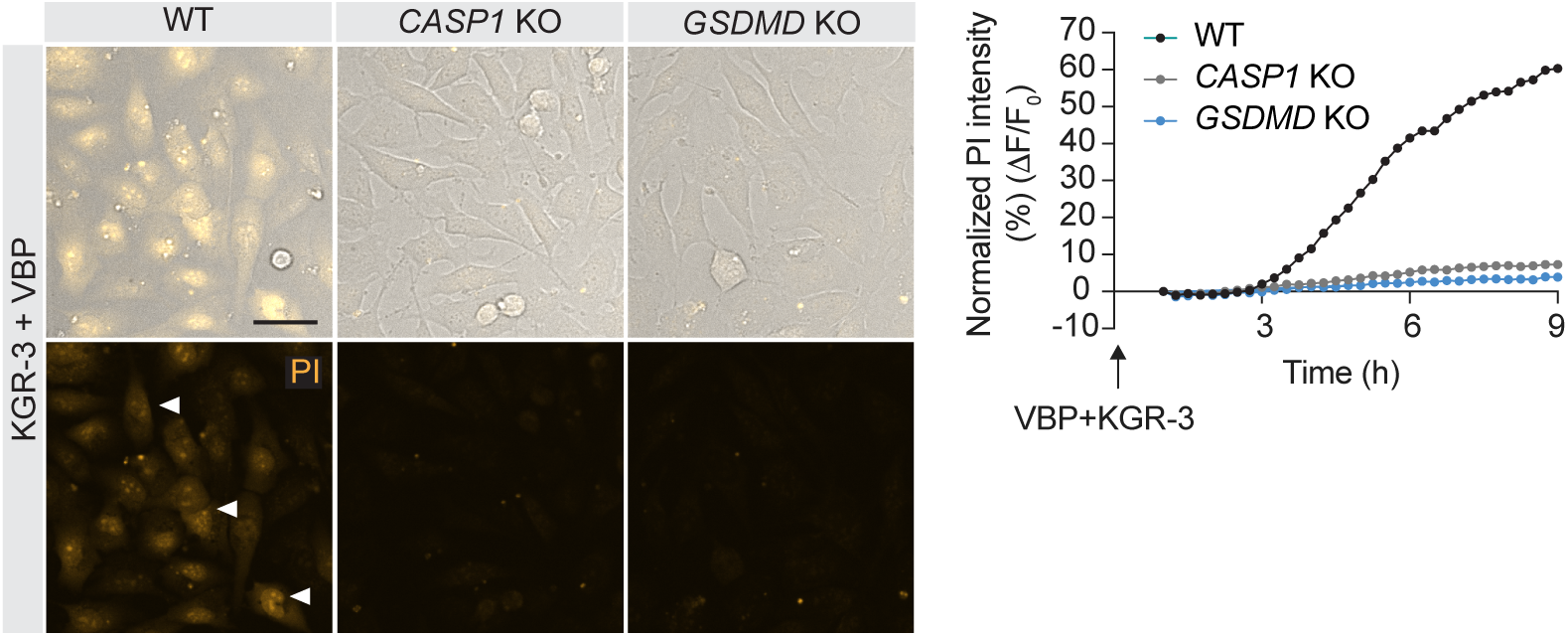
Enhanced membrane permeability in live cells forming GSDMD pores. EA.hy926 cells cultured in 20 μM VBP, 10 μM KGR-3, and 10 μg/ml PI for 6 h. Graph indicates PI fluorescence intensity normalized to the fluorescence at 0 h. Results representative of 3 independent experiments.

## Characterization of inhibitor KGR-53P

Caspase-11-dependent pyroptosis is an important mediator of LPS-induced endotoxic shock in mice ^9,28^, so we also characterized KGR-53, a more potent inhibitor of caspase-11 than KGR-3 (Extended Data Table 2). KGR-53 suppressed but did not prevent caspase-11-dependent pyroptosis in LPS-transfected mouse bone marrow-derived macrophages (BMDMs) (Extended data Fig. 6a). Given that GSDMD pores may transport negatively charged compounds less efficiently ^29^, we derived KGR-53P, which lacks the negative charge of the P3 Glu (Extended Data Fig. 6b). KGR-53P was better than KGR-53 at inhibiting LPS-induced pyroptosis in BMDMs (IC_50_ approximately 4 μM; Fig. 3a, b), despite having very low cell permeability when assessed in healthy gMDCKI cells (Extended Data Table 3). KGR-53P blocked LPS-induced GSDMD cleavage, as well as caspase-11-dependent secretion of IL-1β and IL-18 (Fig. 3c-e). KGR-53P reduced cell death and IL-18 secretion even when added several hours after LPS transfection (Fig. 3e, f). By contrast, KGR-53P failed to protect BMDMs from Venetoclax-induced intrinsic apoptosis and thymocytes from Fas ligand (FasL)-induced extrinsic apoptosis (Extended Data Fig. 6c-g).

**Fig. 3.**
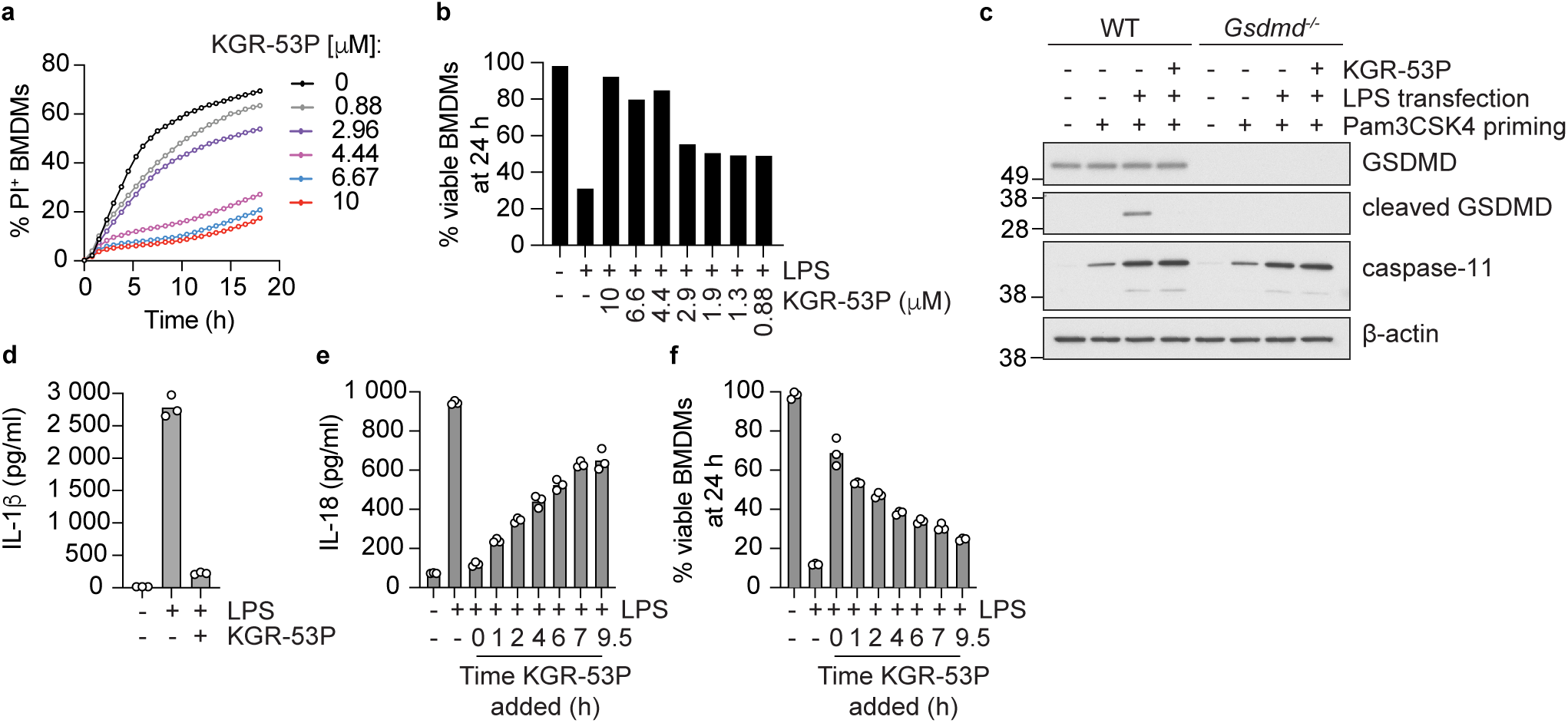
KGR-53P inhibits caspase-11-dependent pyroptosis. **a,** Percentage of BMDMs exhibiting nuclear PI staining after transfection with 5 mg/ml LPS, followed by culture in KGR-53P. Results representative of 2 independent experiments. **b,** Percentage of viable BMDMs in (a) by CellTiter-Glo assay. **c,** Western blots of BMDMs transfected with 5 mg/ml LPS and then cultured in 10 μM KGR-53P for 16 h. Results representative of 2 independent experiments. **d and e,** IL-1β (d) and IL-18 (e) secreted from BMDMs transfected with 5 mg/ml LPS and then cultured in 10 μM KGR-53P for 18 h. In (e), KGR-53P was added at different times after LPS transfection. Bars indicate the mean of cells from different mice (circles, n=3). **f,** Percentage of viable BMDMs in (e) by CellTiter-Glo assay.

Next, we determined whether KGR-53P suppressed caspase-1, caspase-11, and GSDMD-dependent IL-18 and IL-1β production in C57BL/6N mice dosed with LPS from E. Coli O111:B4 ^8,15,24^. As expected, animals receiving 20 mg/kg LPS had significantly elevated IL-18 and IL-1β in their serum after 12 h compared with animals given vehicle alone (Fig. 4). However, markedly less IL-18 and IL-1β was observed when animals were treated with KGR-53P at 30 min before and 6 h after LPS dosing (Fig. 4). Therefore, an inhibitor of caspase-1/11 that is impermeable in healthy cells can nonetheless limit the GSDMD-dependent secretion of proinflammatory IL-1β and IL-18 in vivo, presumably accessing its intracellular target caspases through GSDMD pores. This finding indicates that caspase inhibitors with poor cell permeability profiles may yet have therapeutic potential in the treatment of diseases characterized by excessive inflammasome activation, including sepsis, acute respiratory distress syndrome, and COVID-19 ^30–32^.

**Fig. 4.**
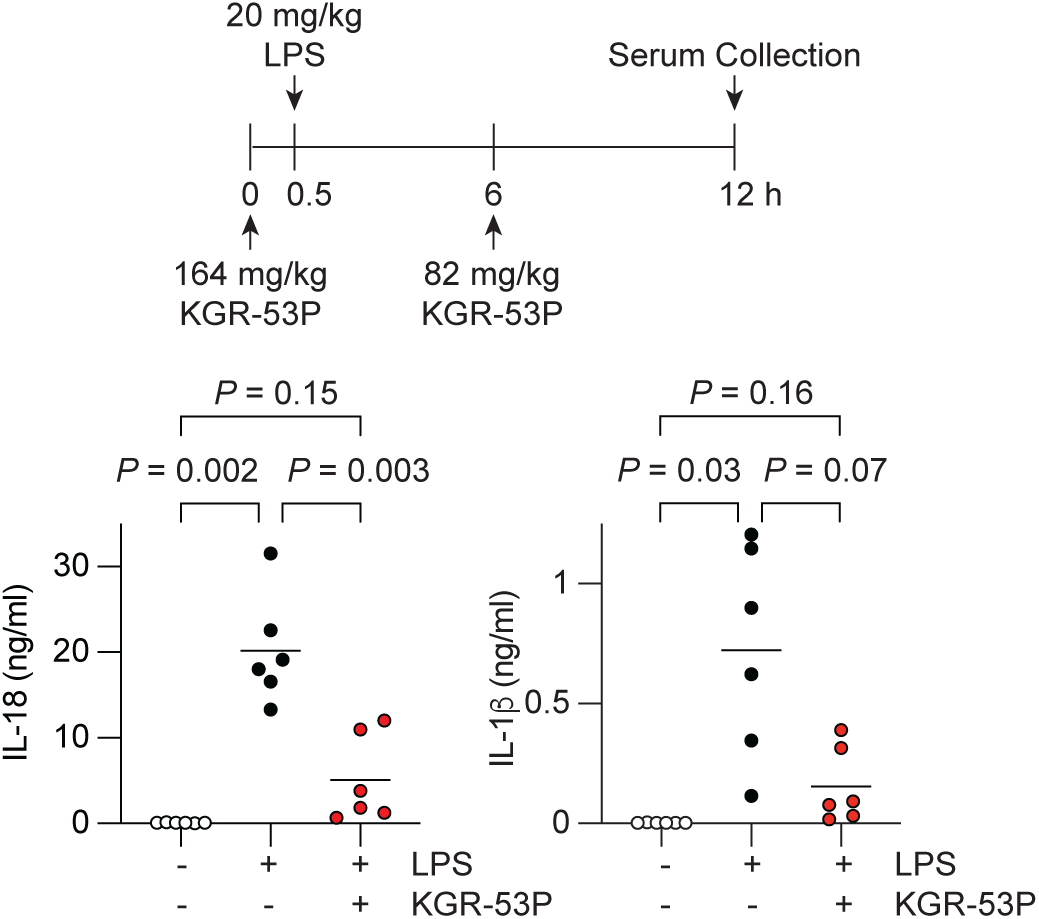
KGR-53P reduces serum IL-1β and IL-18 in mice treated with LPS. Serum levels of IL-1β and IL-18 in female C57BL/6N mice at 12 h after LPS dosing. *P*-values determined by Brown-Forsythe and Welch Anova Tests.

## Methods

### Reagents

All chemicals were from commercial sources and used without further purification. Fluorenylmethyloxycarbonyl (Fmoc)- and tert-butyloxycarbonyl (Boc)-protected amino acids were from Iris Biotech GmbH (Germany), Sigma–Aldrich (Poland), Bachem (USA), Creosalus (USA), and PE Biosciences Limited (Hong Kong). Rink amide AM resin (200-300 mesh, loading 0.74 mmol/g), 2-chlorotrityl chloride resin (100-200 mesh, loading 1.59 mmol/g), biotin, HBTU, HATU, piperidine (PAP), diisopropylcarbodiimide (DICI), 2,2,2-trifluoroethanol (TFE), and trifluoroacetic acid (TFA) were from Iris Biotech GmbH. Anhydrous HOBt was from Creosalus. 2,4,6-Collidine (2,4,6-trimethylpyridine), acetonitrile (ACN, HPLC gradient grade), triisopropylsilane (TIPS), hydrobromic acid solution (30% HBr wt. in acetic acid), *N*-methylmorpholine (NMM), tetrahydrofuran (THF, anhydrous), isobutylchloroformate (IBCF), and 2,6-dimethylbenzoic acid (2,6-DMBA) were from Sigma–Aldrich. N,N’-Dimethylformamide (DMF, pure for analysis), methanol (MeOH), dichloromethane (DCM), acetic acid (AcOH), diethyl ether (Et_2_O), and phosphorus pentoxide (P_2_O_5_) were from POCh (Poland). Diazomethane used for the synthesis of AOMK inhibitors was generated according to the Aldrich Technical Bulletin (AL-180) protocol. All compounds (peptides, ACC fluorescent substrates, AIE fluorescent substrates, and inhibitors) were purified by reverse-phase HPLC on a Waters system (Waters M600 solvent delivery module and Waters M2489 detector system) using a semipreparative Discovery C8 column (particle size 10 μm). Compound purity was confirmed by HPLC using an analytical Discovery C8 column (particle size 10 μm). Phase A was water/0.1% TFA. Phase B was ACN/0.1% TFA. For purification and compound analysis, the assay was run for 30 min in a linear gradient (from 5% phase B to 100% phase B). The molecular weight of each synthesized compound was confirmed on a WATERS LCT Premier XE High Resolution Mass Spectrometer with electrospray ionization (ESI) and a time of flight (TOF) module.

### Synthesis of caspase substrates

Substrates were synthesized as described ^6^. In brief, Fmoc-P2-OH (2.5 eq) was preactivated with HATU (2.5 eq) and 2,4,6-collidine (2.5 eq) in DMF, then added to a cartridge containing H_2_N-Asp(O-tBu)-ACC-resin. The mixture was gently agitated for 3 h, filtered, and washed with DMF. The Fmoc-protecting group was subsequently removed using 20% piperidine in DMF, with a ninhydrin test performed after each coupling and deprotection step. A solution of Fmoc-P3-OH (2.5 eq.), HATU (2.5 eq.), and 2,4,6-collidine (2.5 eq.) in DMF was added to the resin, followed by agitation for 3 h. Next, the Fmoc group was deprotected, and Fmoc-P4-OH amino acid was conjugated in a similar manner. After Fmoc deprotection from the P4 position, the N-terminus was protected with an acetyl group using AcOH (5 eq.), HBTU (5 eq.), and DIPEA (5 eq.) in DMF. Following solvent removal, the resin was washed with DMF DCM, and MeOH, then dried over P_2_O_5_ and cleaved from the resin with a mixture of TFA/TIPS/H_2_O. The crude product was purified by HPLC and lyophilized. Purity and molecular weight were confirmed with LC-MS. Substrates were dissolved in peptide-grade DMSO to a concentration of 20 mM and stored at −80°C.

### Synthesis of caspase inhibitors

Ac-P4-P3-P2-P1-AOMK inhibitors were synthesized as described ^34,35^. In brief, an Ac-P4-P3-P2-COOH peptide was synthesized on a solid support (2-chlorotrityl chloride resin) and utilized for further synthesis without purification. Simultaneously, Boc-Asp(O-tBu)-AOMK was generated through the reaction of Boc-Asp(O-tBu)-COOH with diazomethane, followed by conversion to bromomethyl ketone (-CH_2_Br) and subsequently to 2,6-dimethyl acyloxymethyl ketone (-AOMK). Next, Boc-Asp(O-tBu)-AOMK was deprotected with TFA to yield H_2_N-Asp-AOMK. The N-deprotected warhead was then coupled to Ac-P4-P3-P2-COOH to obtain Ac-P4-P3-P2-Asp-AOMK. Following removal of the protecting groups from P4-P2 amino acids using TFA, the final crude product was purified by RP-HPLC on a C8 column.

### Recombinant caspases and kinetic assays

Caspases were expressed, purified, and active site titrated using z-VAD-FMK (Promega) as described ^36^. Kinetic assays were conducted at 37°C using an fMax fluorimeter (Molecular Devices) operating in kinetic mode. Excitation for ACC-based substrates was at 355 nm, while emission was detected at 460 nm with a cutoff set at 455 nm. The assay buffer was 1M sodium citrate, 10% w/v sucrose, 20 mM Pipes, 10 mM NaCl, 1 mM EDTA, and 10 mM DTT, pH 7.2– 7.4, except for caspase-3 and -7, when 1M sodium citrate was omitted. In Extended Data Fig. 1, caspases were incubated at 37°C for 15 min prior to mixing with 100 μM substrate in 100 μl. Reactions contained 3 nM caspase-1, 100 nM caspase-4, or 120 nM caspase-5, and were monitored for 30 min. Alternatively, E. coli expressing one of the caspases were resuspended in 50 mM Tris pH 7.4, 100 mM NaCl and sonicated for 2 min at full power with 50% duty cycle (on for 0.5 s, then off for 0.5 s, Branson Sonifier 250). Lysates were centrifuged for 30 min at 18,000 x g. Supernatants were filtered (0.45 μm) and diluted two-fold with 2x buffer (described above) prior to measurements. The total protein concentration in the lysates varied between 150-300 mg/ml, and the final concentration of the lysate was adjusted during caspase activity measurements with an appropriate substrate (e.g., Ac-DEVD-ACC for caspase-3, Ac-WEHD-ACC for caspase-1, -4, -5, etc.). Hybrid Combinatorial Substrate Libraries were screened 100 μM substrate. Velocity calculations were performed using the linear phase of the progress curve, typically observed between 15–30 min. Experiments were performed in triplicate, and reported results are mean values, with variation between individual measurements consistently <10%. Data analysis involved normalizing total Relative Fluorescence Units (RFUs) for each sub-library, with the highest value set to 100% and adjustments made accordingly.

In Extended Data Table 1, reactions contained 2-15 nM caspase-1, 3-35 nM caspase-3, 35-200 nM caspase-4, 20-250 nM caspase-5, 40-400 nM caspase-7, 6-42 nM caspase-8, or 20-150 nM caspase-9, and were performed using eight different substrate concentrations between 0.05 and 500 μM. DMSO in the reactions was kept below 2% (v/v). Each reaction was done in triplicate. k_cat/_K_M_ calculations were performed using GraphPad Prism version 8.0.2.

In Extended Data Table 2, the k_obs_/I parameter, which represents the pseudo first-order kinetic constant, was determined using the equation: Y = (V_0_ − V_S_) × (1 − exp(−k_obs_[app])) / k_obs_[app] + V_S_ × X, where Y denotes enzyme activity, V_0_ stands for initial enzyme activity, and V_S_ represents enzyme activity at equilibrium. Initially, we screened all inhibitors in triplicate at concentrations of 10 nM, 100 nM, 1 μM, and 10 μM against each caspase. Subsequently, we calculated the k_obs_[app] /I values from the plots that exhibited the best fitting to the equation curve. The k_obs_[app] /I values for the most effective inhibitors were derived from the analyses conducted at 10 nM and 100 nM inhibitor, while those for less effective inhibitors were determined using 1 μM and 10 μM inhibitor. We calculated k_obs_/I using the formula k_obs_/I = k_obs_/I*(1+K_M_/[S]), where K_M_ represents the Michaelis-Menten constant for the substrate used, and [S] denotes the substrate concentration. Each caspase was incubated with an appropriate ACC-labeled substrate to monitor the reaction: caspase-1, Ac-WEHD-ACC, [S] = 50 μM, K_M_ = 11 μM; caspase-3, Ac-DEVD-ACC, [S] = 50 μM, K_M_ = 20 μM; caspase-4, Ac-WEHD-ACC, [S] = 100 μM, K_M_ = 43 μM; caspase-6, Ac-VEID-ACC, [S] = 100 μM, K_M_ = 42 μM; caspase-7, Ac-DEVD-ACC, [S] = 100 μM, K_M_ = 48 μM; caspase-8, Ac-LEHD-ACC, [S] = 50 μM, K_M_ = 8 μM; caspase-9, Ac-LEHD-ACC, [S] = 200 μM, K_M_ = 102 μM; caspase-10, Ac-LEHD-ACC, [S] = 50 μM, K_M_ = 27 μM.

In Extended Data Table 3, inhibitors were examined with greater rigor against both human and mouse caspases. Mouse caspase-1, Ac-WEHD-ACC, [S] = 50 μM, K_M_ = 12.5 μM; mouse caspase-1, Ac-DEVD-ACC, [S] = 50 μM, K_M_ = 25 μM; mouse caspase-8, Ac-LEHD-ACC, [S] = 50 μM, K_M_ = 6.5 μM; mouse caspase-11, Ac-WEHD-ACC, [S] = 200 μM, K_M_ = 132 μM. Reactions contained 1.9 nM human caspase-1, 2.3 nM human caspase-3, 43 nM human caspase-4, 4.1 nM human caspase-8, 14 nM human caspase-9, 2.2 nM mouse caspase-1, 1.8 nM mouse caspase-3, 3.2 nM mouse caspase-8, or 38 nM mouse caspase-11, ensuring that the lowest inhibitor concentration was at least 4-fold higher than the caspase concentration. Inhibitors were diluted in 384-well plates, mixed with substrate, and preincubated at 37°C. Caspases were preincubated separately at 37°C for 15 min, and then added to the mix of inhibitor and substrate. Fluorescence was monitored for 20 min. K_obs_ values were calculated as before, and k_obs_/I was determined by plotting k_obs_ values against [I] and applying linear regression. Parameters were derived from at least 3 independent experiments and presented as average values.

### Cells

THP-1 cells (ATCC; mycoplasma tested, but not authenticated) were maintained in RPMI-1640 medium supplemented with 2 mM L-glutamine, 10% fetal bovine serum (FBS; Omega Scientific), 100 U/ml penicillin, and 100 μg/ml streptomycin (Gibco). EA.hy926 cells (ATCC; mycoplasma tested, but not authenticated) were grown in the high glucose version of Dulbecco’s modified Eagle’s medium (DMEM) supplemented with 10 mM HEPES pH 7.4, 1x GlutaMAX (Gibco), 100 U/ml penicillin, and 100 μg/ml streptomycin, 1x non-essential amino acids (Gibco), 1 mM sodium pyruvate (Gibco), and 10% FBS.

To generate *GSDMD* KO cells, THP-1 cells stably expressing Cas9 were electroporated with sgRNAs (IDT) using the Neon electroporation system (ThermoFisher). Target sequences in human *GSDMD* were CACAAGCGTTCCACGAGCGA and ACGCGCACCCACAAGCGGGA. To generate Ea.hy926 *CASP1* or *GSDMD* KO cells, Cas9+ cells were transduced with gRNAs (CCTCAAACTCTTCTGTAGT for *CASP1*) by lentiviral delivery (pLKO.1 vector, Sigma), followed by selection of gRNA-expressing cells with 2 μg/ml puromycin (Gibco). To express FKBP-Casp4 and FKBP-Casp11 fusions, the dimerizer domain (DmrB) of a modified FK506 binding protein (FKBP) was added as an N-terminal fusion to the catalytic domains of caspase-4 (aa 88-377) and caspase-11 (aa 88-373). cDNAs encoding the fusion proteins were generated in a lentiviral vector for EA.hy926 cells or piggyBac vector BH1.11 (Genentech) for THP-1 cells. After lentiviral transduction, EA.hy926 cells were selected with 2 μg/ml puromycin (Gibco). THP-1 cells expressing Cas9 were co-electroporated with FKBP-Casp4/BH1.11 or FKBP-Casp11/BH1.11 and the transposase vector pBO (Transposagen Biopharmaceuticals) using the Neon system. THP-1 cells were selected with 10 μg/ml Blasticidin S HCl (Gibco). THP-1 and EA.hy926 cells were treated with 50 nM and 2 nM B/B homodimerizer (AP20187, MedChemExpress), respectively, to activate the FKBP-caspase fusions.

Bone marrow cells from C57BL/6N mice were differentiated into BMDMs in DMEM supplemented with 10% FBS, 1x GlutaMAX, 100 U/ml penicillin, 100 μg/ml streptomycin, and 30% L929-conditioned medium for 5-6 days. Thymocytes from C57BL/6N mice were cultured in the high glucose version of DMEM supplemented with 10 mM HEPES pH 7.4, 1x GlutaMAX (Gibco), 100 U/ml penicillin, 100 μg/ml streptomycin, 1x non-essential amino acids, 50 μM 2-mercaptoethanol, and 10% FBS.

### *In vitro* Permeability Assays

Permeability assays in gMDCKI cells (mycoplasma tested, but not authenticated) were performed as described ^13^. Briefly, cells were seeded at 7×10^5^ cells/ml in 96-well transwells in 100 μl DMEM containing phenol red, 10% BSA, 1% pen/strep, 1x GlutaMAX, and 5µg/ml plasmocin, grown until confluent (around 4 days), and then placed in Hank’s Balanced Salt Solution containing 25 mM sterile filtered HEPES buffer, pH 7.4 (Invitrogen, Grand Island, NY, USA). 25 µM caspase inhibitor was added to the donor compartment together with the monolayer integrity marker Lucifer Yellow (Sigma Aldrich, St. Louis, MO, USA). Inhibitor concentrations in the donor and receiver compartments after 3 h at 37°C were determined by LC-MS/MS. The apparent permeability (P_app_) of test compounds was determined using the equation P_app_ = (dQ/dt)*(1/AC_0_), where dQ/dt is the rate of compound appearance in the receiver compartment, Q is the quantity of compound, C_0_ is the concentration in the donor compartment and A is the surface area of the insert. Efflux ratio was calculated as (Papp, B-to-A)/(Papp, A-to-B), where A is apical and B is basolateral.

### Cell Death and Cytokine Assays

THP-1 cells were seeded in Opti-MEM (Gibco) supplemented with 0.1% FBS, primed for 4 h with 1 μg/ml ultrapure LPS (*Escherichia coli* O111:B4; InvivoGen), and then stimulated for 2 h with 5 μM nigericin (InvivoGen) ± 25 μM emricasan (Abcam), 25 μM tetrapeptide-AOMK, or 25 μM MCC950 (InvivoGen). THP-1 cells were treated for 24 h with 1 µM staurosporine ± 10 µM emricasan or tetrapeptide-AOMK. THP-1 cells expressing FKBP-caspase-4 or -11 were treated for 24 h with 50 nM B/B ± 10µM emricasan or tetrapeptide-AOMK. LDH release into the culture medium was measured with a CytoTox 96 Non-Radioactive Cytotoxicity Assay (Promega). Results are calculated as a percentage of maximum LDH release by 0.1% Triton-X100. Cell viability was determined using a CellTiter-Glo Luminescent Cell Viability Assay (Promega).

EA.hy926 cells (8,000-10,000 per well) were seeded in 96-well plates overnight and then stimulated in Opti-MEM containing 0.1% FBS with either 20 μM Valboro-Pro (MilliporeSigma) or 100 ng/ml TRAIL (ALX-201-073-C020; Enzo Life Sciences) ± 10 μM tetrapeptide-AOMK. Viability was measured by CellTiter Glo assay.

BMDMs (100,000 per well) were plated in 96-well plates overnight. For inflammasome stimulations, cells were primed with 1 μg/ml Pam3CSK4 (InvivoGen) for 5 h to induce expression of *Casp11*. Primed cells were transfected with 5 μg/ml LPS using Fugene-HD transfection reagent (Promega). Unprimed BMDMs were stimulated with 25 μM Venetoclax (R&D Systems). Where indicated, BMDMs were treated with 10 µM tetrapeptide-AOMK. IL-1β in culture supernatants was measured using a 384-well multi-array mouse IL-1β assay (Meso Scale Diagnostics). IL-18 secretion was determined using a mouse IL-18 ELISA kit (MBL International Corporation).

Mouse thymocytes were stimulated with 1 μg/ml FasL (Adipogen) ± 10 μM emricasan or KGR-53P. Cells were stained with 2.5 μg/ml propidium iodide (PI; BD Biosciences) and analysed in a FACSCanto II cytometer (BD Biosciences) with BD FACSDiva8.0. Viable cells were PI-negative.

### Western blots

In Extended Data Fig. 3a, cells were lysed in 10 mM Tris-HCl pH 7.5, 150 mM NaCl, 2.5 mM MgCl2, 0.5 mM CaCl_2_, 1% NP40, phosphatase and protease inhibitors (Roche), and DNase (approximately 80 U/ml, Qiagen). For all other western blots, cells were lysed in 20 mM Tris-HCl pH 7.5, 135 mM NaCl, 1.5 mM MgCl_2_, 10% glycerol, 1 mM EGTA, and 1% Triton X-100 supplemented with protease and phosphatase inhibitors (Roche). The soluble lysate was recovered after centrifugation for 15 min at 20,000g and 4°C. Westerns blots used antibodies recognising human and mouse GSDMD (AB219800, Abcam), caspase-3 (9662, Cell Signaling Technology), cleaved caspase-3 (9664, Cell Signaling Technology), caspase-4 (4B9, Enzo Life Sciences), caspase-7 (9492, Cell Signaling Technology), cleaved caspase-7 (9491, Cell Signaling Technology), caspase-8 (4927, Cell Signaling Technology), cleaved caspase-8 (8592, Cell Signaling Technology), caspase-9 (9508, Cell Signaling Technology), caspase-11 (NB-120-10454, Novus), FKBP12 (ab2918, Abcam), HRP-conjugated GAPDH (3683, Cell Signaling Technology), HRP-conjugated β-actin (5125S, Cell Signaling Technology) or α-tubulin (66031-1-Ig, Proteintech). HRP-conjugated secondary antibodies were purchased from Jackson Immunoresearch.

### Live cell imaging

EA.hy926 cells (3×10^4^ per well) were plated in 8-well chamber slides (Ibidi #80806). Brightfield and fluorescence images were acquired every 15 min using a Nikon TI-E Spinning Disk Confocal equipped with a 20x Apo 0.75 NA DIC, WD1 objective (Nikon) and a Photometrics Prime95B sCMOS camera (Teledyne Photometrics). Samples were maintained at 37°C and 5% CO_2_ throughout imaging using an environmental control chamber and gas mixer (Oko Labs). Cells were treated in Opti-MEM containing 10 mg/ml PI. Images were processed in the Nikon Elements software or Fiji. For quantification of cytoplasmic PI entry, cells were segmented using the Trackmate plugin on Fiji ^37^. Fluorescence intensity was measured after thresholding and background fluorescence subtraction. Relative intensity change was calculated using the fluorescence intensity ratio F/F_0_, where F_0_ is the cellular fluorescence at time zero.

Plasma membrane integrity was assessed by culturing cells in 1 μg/ml PI (MilliporeSigma) in an IncuCyte S3 (Essen BioScience). Incubation with 1% Triton X-100 was used to record the total number of cells at the final timepoint. Data are presented as the percentage of total cells that have PI+ nuclei. For representative images of PI-positive cells, cells were plated on a glass-bottom 96 well plate (655892, Greiner Bio-One) and imaged with a 20x objective on the IncuCyte S3.

### Mice

All mouse studies complied with relevant ethics regulations and were approved by the Genentech Institutional Animal Care and Use Committee (IACUC) in an Association for Assessment and Accreditation of Laboratory Animal Care (AAALAC)-accredited facility. Female C57BL/6N mice aged 9 weeks (Charles River Labs) were dosed intraperitoneally with 164 mg/kg KGR-53P in vehicle (20 mM phosphate pH 7 with 3.75% Solutol HS 15) or vehicle alone at 0 h, 20 mg/kg LPS (E. Coli O111:B4, Sigma Aldrich) in PBS or PBS alone at 30 min, and 82 mg/kg KGR-53P or vehicle alone at 6 h. Serum IL-18 after 12 h was determined by mouse IL-18 ELISA (7625, MBL International Corporation). Serum IL-1β was determined by the V-PLEX Plus Mouse IL-1β kit (K152QPG, Meso Scale Diagnostics).

## Acknowledgments

We thank Dixit Lab members, Mikiko Okumura, Michael Koehler, Sarah Gierke, Praveen Krishnamoorthy, and members of the Genentech Center for Advanced Light Microscopy for discussions, reagents, and technical assistance. We thank Scott Snipas and Guy Salvesen for recombinant caspases. M.P. is supported by the National Science Centre in Poland (OPUS grant, UMO-2018/29/B/NZ1/02249).

## Author Contributions

The work reported herein was primarily carried out by K. M. G. as a participant in the Genentech Postoctoral Program (https://careers.gene.com/us/en/students-postdocs); The caspase inhibitors were designed at Wroclaw University of Science and Technology by K. M. G. under the supervision of M. P.; K. M. G., J. N. and M. K. synthesized inhibitors as part of the HyCoSuL screening library, a method developed by M. D. ^33^; K. M. G., J. N. and M. K. performed kinetic analyses under the supervision of M. P.; K. M. G., M. E. T., R. S. J., N. K., K. N., M. P. and V. M. D. contributed to the design of experiments; K. M. G. and M. E. T. performed cell culture experiments; I. S. and B. L generated cell lines; E. S. L. and P. K. optimized inhibitor formulations; W. P. L. and J. Zhang performed mouse studies; K. N. wrote this paper with input from all co-authors.

## Competing interests

K.M.G., M.E.T., I.S., B.L., R.S.J., E.S.L., P.K., W.P.L, J.Z., N.K., K.N., and V.M.D. were employees of Genentech.

Correspondence and requests for materials should be addressed to V.M.D.

---

**Extended Data Table 1.**
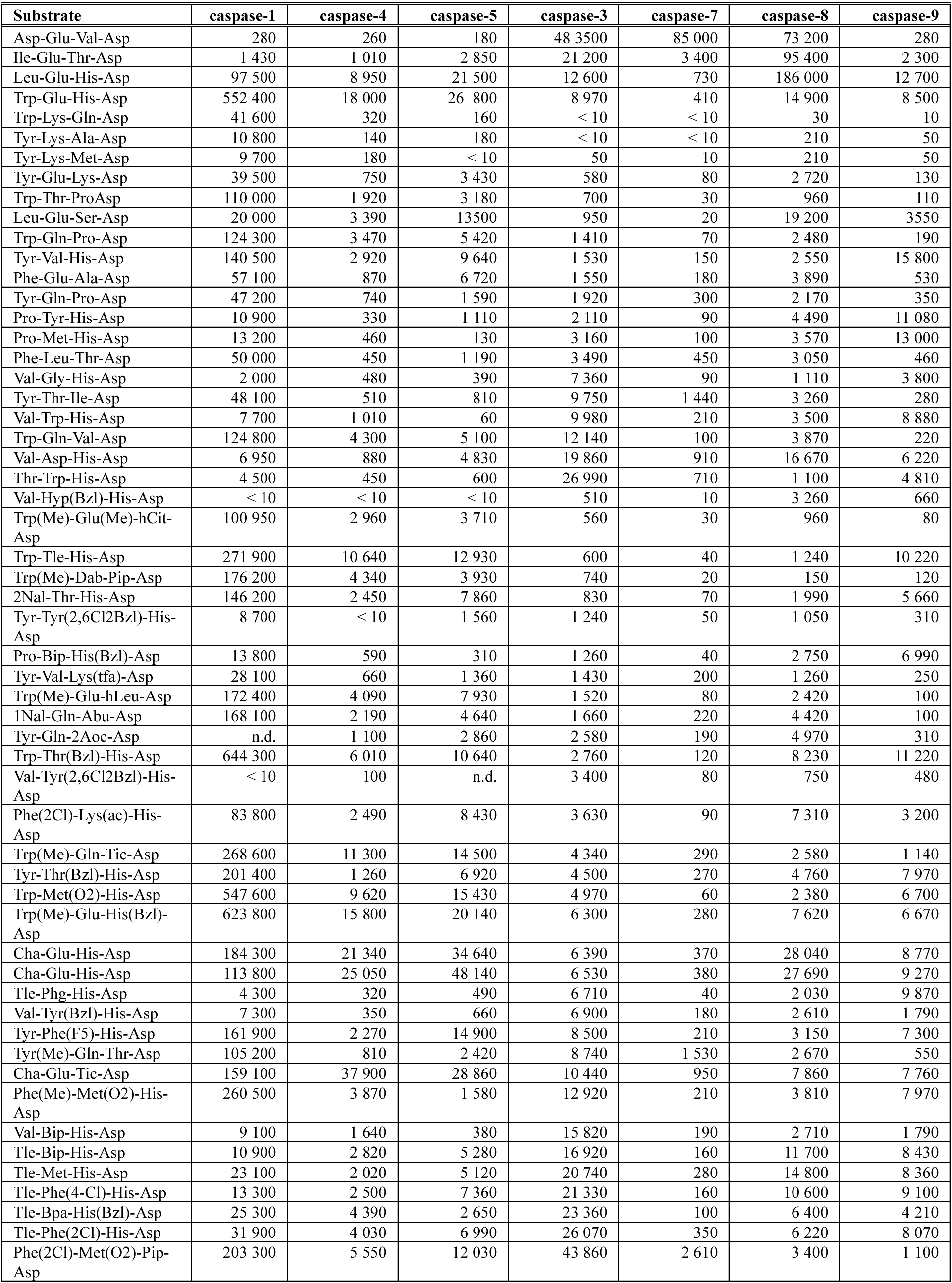
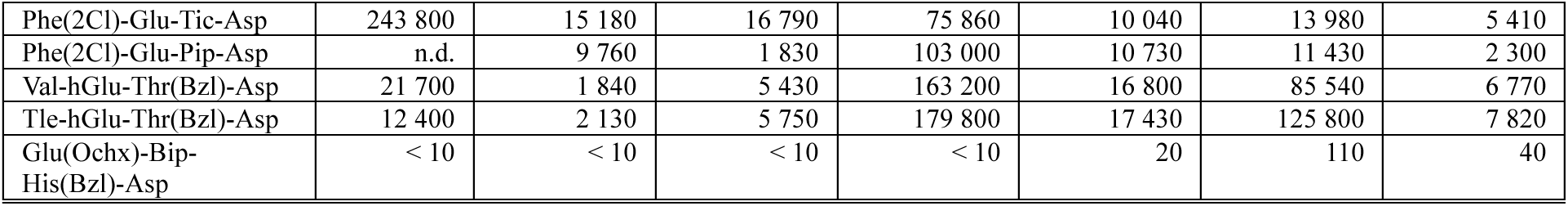
Human caspase k_cat/_K_M_ values (M^-1^s^-1^) for tetrapeptide substrates having the format Ac (acetyl)-P4-P3-P2-P1(Asp)-ACC (7-amino-4-carbamoylmethylcoumarin). n.d., not determined. See methods for the details.

| Substrate | caspase-1 | caspase-4 | caspase-5 | caspase-3 | caspase-7 | caspase-8 | caspase-9 |
| --- | --- | --- | --- | --- | --- | --- | --- |
| Asp-Glu-Val-Asp | 280 | 260 | 180 | 48 3500 | 85 000 | 73 200 | 280 |
| Ile-Glu-Thr-Asp | 1 430 | 1 010 | 2 850 | 21 200 | 3 400 | 95 400 | 2 300 |
| Leu-Glu-His-Asp | 97 500 | 8 950 | 21 500 | 12 600 | 730 | 186 000 | 12 700 |
| Trp-Glu-His-Asp | 552 400 | 18 000 | 26 800 | 8 970 | 410 | 14 900 | 8 500 |
| Trp-Lys-Gln-Asp | 41 600 | 320 | 160 | < 10 | < 10 | 30 | 10 |
| Tyr-Lys-Ala-Asp | 10 800 | 140 | 180 | < 10 | < 10 | 210 | 50 |
| Tyr-Lys-Met-Asp | 9 700 | 180 | < 10 | 50 | 10 | 210 | 50 |
| Tyr-Glu-Lys-Asp | 39 500 | 750 | 3 430 | 580 | 80 | 2 720 | 130 |
| Trp-Thr-ProAsp | 110 000 | 1 920 | 3 180 | 700 | 30 | 960 | 110 |
| Leu-Glu-Ser-Asp | 20 000 | 3 390 | 13500 | 950 | 20 | 19 200 | 3550 |
| Trp-Gln-Pro-Asp | 124 300 | 3 470 | 5 420 | 1 410 | 70 | 2 480 | 190 |
| Tyr-Val-His-Asp | 140 500 | 2 920 | 9 640 | 1 530 | 150 | 2 550 | 15 800 |
| Phe-Glu-Ala-Asp | 57 100 | 870 | 6 720 | 1 550 | 180 | 3 890 | 530 |
| Tyr-Gln-Pro-Asp | 47 200 | 740 | 1 590 | 1 920 | 300 | 2 170 | 350 |
| Pro-Tyr-His-Asp | 10 900 | 330 | 1 110 | 2 110 | 90 | 4 490 | 11 080 |
| Pro-Met-His-Asp | 13 200 | 460 | 130 | 3 160 | 100 | 3 570 | 13 000 |
| Phe-Leu-Thr-Asp | 50 000 | 450 | 1 190 | 3 490 | 450 | 3 050 | 460 |
| Val-Gly-His-Asp | 2 000 | 480 | 390 | 7 360 | 90 | 1 110 | 3 800 |
| Tyr-Thr-Ile-Asp | 48 100 | 510 | 810 | 9 750 | 1 440 | 3 260 | 280 |
| Val-Trp-His-Asp | 7 700 | 1 010 | 60 | 9 980 | 210 | 3 500 | 8 880 |
| Trp-Gln-Val-Asp | 124 800 | 4 300 | 5 100 | 12 140 | 100 | 3 870 | 220 |
| Val-Asp-His-Asp | 6 950 | 880 | 4 830 | 19 860 | 910 | 16 670 | 6 220 |
| Thr-Trp-His-Asp | 4 500 | 450 | 600 | 26 990 | 710 | 1 100 | 4 810 |
| Val-Hyp(Bzl)-His-Asp | < 10 | < 10 | < 10 | 510 | 10 | 3 260 | 660 |
| Trp(Me)-Glu(Me)-hCit-Asp | 100 950 | 2 960 | 3 710 | 560 | 30 | 960 | 80 |
| Trp-Tle-His-Asp | 271 900 | 10 640 | 12 930 | 600 | 40 | 1 240 | 10 220 |
| Trp(Me)-Dab-Pip-Asp | 176 200 | 4 340 | 3 930 | 740 | 20 | 150 | 120 |
| 2Nal-Thr-His-Asp | 146 200 | 2 450 | 7 860 | 830 | 70 | 1 990 | 5 660 |
| Tyr-Tyr(2,6Cl2Bzl)-His-Asp | 8 700 | < 10 | 1 560 | 1 240 | 50 | 1 050 | 310 |
| Pro-Bip-His(Bzl)-Asp | 13 800 | 590 | 310 | 1 260 | 40 | 2 750 | 6 990 |
| Tyr-Val-Lys(tfa)-Asp | 28 100 | 660 | 1 360 | 1 430 | 200 | 1 260 | 250 |
| Trp(Me)-Glu-hLeu-Asp | 172 400 | 4 090 | 7 930 | 1 520 | 80 | 2 420 | 100 |
| 1Nal-Gln-Abu-Asp | 168 100 | 2 190 | 4 640 | 1 660 | 220 | 4 420 | 100 |
| Tyr-Gln-2Aoc-Asp | n.d. | 1 100 | 2 860 | 2 580 | 190 | 4 970 | 310 |
| Trp-Thr(Bzl)-His-Asp | 644 300 | 6 010 | 10 640 | 2 760 | 120 | 8 230 | 11 220 |
| Val-Tyr(2,6Cl2Bzl)-His-Asp | < 10 | 100 | n.d. | 3 400 | 80 | 750 | 480 |
| Phe(2Cl)-Lys(ac)-His-Asp | 83 800 | 2 490 | 8 430 | 3 630 | 90 | 7 310 | 3 200 |
| Trp(Me)-Gln-Tic-Asp | 268 600 | 11 300 | 14 500 | 4 340 | 290 | 2 580 | 1 140 |
| Tyr-Thr(Bzl)-His-Asp | 201 400 | 1 260 | 6 920 | 4 500 | 270 | 4 760 | 7 970 |
| Trp-Met(O2)-His-Asp | 547 600 | 9 620 | 15 430 | 4 970 | 60 | 2 380 | 6 700 |
| Trp(Me)-Glu-His(Bzl)-Asp | 623 800 | 15 800 | 20 140 | 6 300 | 280 | 7 620 | 6 670 |
| Cha-Glu-His-Asp | 184 300 | 21 340 | 34 640 | 6 390 | 370 | 28 040 | 8 770 |
| Cha-Glu-His-Asp | 113 800 | 25 050 | 48 140 | 6 530 | 380 | 27 690 | 9 270 |
| Tle-Phg-His-Asp | 4 300 | 320 | 490 | 6 710 | 40 | 2 030 | 9 870 |
| Val-Tyr(Bzl)-His-Asp | 7 300 | 350 | 660 | 6 900 | 180 | 2 610 | 1 790 |
| Tyr-Phe(F5)-His-Asp | 161 900 | 2 270 | 14 900 | 8 500 | 210 | 3 150 | 7 300 |
| Tyr(Me)-Gln-Thr-Asp | 105 200 | 810 | 2 420 | 8 740 | 1 530 | 2 670 | 550 |
| Cha-Glu-Tic-Asp | 159 100 | 37 900 | 28 860 | 10 440 | 950 | 7 860 | 7 760 |
| Phe(Me)-Met(O2)-His-Asp | 260 500 | 3 870 | 1 580 | 12 920 | 210 | 3 810 | 7 970 |
| Val-Bip-His-Asp | 9 100 | 1 640 | 380 | 15 820 | 190 | 2 710 | 1 790 |
| Tle-Bip-His-Asp | 10 900 | 2 820 | 5 280 | 16 920 | 160 | 11 700 | 8 430 |
| Tle-Met-His-Asp | 23 100 | 2 020 | 5 120 | 20 740 | 280 | 14 800 | 8 360 |
| Tle-Phe(4-Cl)-His-Asp | 13 300 | 2 500 | 7 360 | 21 330 | 160 | 10 600 | 9 100 |
| Tle-Bpa-His(Bzl)-Asp | 25 300 | 4 390 | 2 650 | 23 360 | 100 | 6 400 | 4 210 |
| Tle-Phe(2Cl)-His-Asp | 31 900 | 4 030 | 6 990 | 26 070 | 350 | 6 220 | 8 070 |
| Phe(2Cl)-Met(O2)-Pip-Asp | 203 300 | 5 550 | 12 030 | 43 860 | 2 610 | 3 400 | 1 100 |
| Phe(2Cl)-Glu-Tic-Asp | 243 800 | 15 180 | 16 790 | 75 860 | 10 040 | 13 980 | 5 410 |
| Phe(2Cl)-Glu-Pip-Asp | n.d. | 9 760 | 1 830 | 103 000 | 10 730 | 11 430 | 2 300 |
| Val-hGlu-Thr(Bzl)-Asp | 21 700 | 1 840 | 5 430 | 163 200 | 16 800 | 85 540 | 6 770 |
| Tle-hGlu-Thr(Bzl)-Asp | 12 400 | 2 130 | 5 750 | 179 800 | 17 430 | 125 800 | 7 820 |
| Glu(Ochx)-Bip-His(Bzl)-Asp | < 10 | < 10 | < 10 | < 10 | 20 | 110 | 40 |

**Extended Data Table 2.** Human caspase k_obs_/I values (M^-1^s^-1^) for tetrapeptide inhibitors having the format Ac-P4-P3-P2-P1(Asp)-AOMK. See methods for how this parameter was calculated.

| Inhibitor | Peptide sequence | caspase-1 | caspase-4 | caspase-3 | caspase-6 | caspase-7 | caspase-8 | caspase-9 | caspase-10 |
| --- | --- | --- | --- | --- | --- | --- | --- | --- | --- |
| KGR-1 | Asp-Glu-Val-Asp | 72 300 | 21 000 | 3 704 000 | 330 000 | 549 200 | 889 000 | 23 100 | 183 000 |
| KGR-2 | Ile-Glu-Thr-Asp | 94 500 | 13 000 | 106 000 | 642 000 | 26 300 | 1 144 000 | 56 200 | 255 000 |
| KGR-3 | Trp-Glu-His-Asp | 3 920 000 | 51 800 | 50 200 | 24 700 | 13 300 | 1 330 000 | 58 300 | 107 000 |
| KGR-4 | Leu-Glu-His-Asp | 750 000 | 30 200 | 108 000 | 102 000 | 15 500 | 1 564 000 | 73 900 | 370 000 |
| KGR-5 | Leu-Glu-Thr-Asp | 651 000 | 37 400 | 397 000 | 180 000 | 37 500 | 1 476 000 | 79 300 | 307 000 |
| KGR-6 | Asp-Glu-Thr(Bzl)-Asp | 23 800 | 8 200 | 4 240 000 | 206 000 | 678 000 | 886 000 | 8 200 | 167 000 |
| KGR-7 | Leu-Glu-Thr(Bzl)-Asp | 54 600 | 39 400 | 345 000 | 351 000 | 51 200 | 628 000 | 18 300 | 159 000 |
| KGR-8 | Leu-Tle-His-Asp | 2 036 000 | 21 500 | 6 700 | 1 300 | 600 | 978 000 | 37 800 | 321 000 |
| KGR-9 | Leu-Tle-Ala-Asp | 3 350 000 | 66 800 | 24 500 | 2 100 | 3 600 | 2 280 000 | 241 600 | 350 000 |
| KGR-10 | Leu-Glu-Ser(Bzl)-Asp | 896 000 | 67 200 | 352 200 | 351 000 | 34 600 | 1 302 000 | 127 200 | 404 000 |
| KGR-13 | Oic-Tle-His-Asp | 6 000 | 1 100 | 5 500 | 900 | 700 | 187 000 | 14 900 | 4 300 |
| KGR-14 | Leu-Glu-hCha-Asp | 307 000 | 38 000 | 122 000 | 493 000 | 17 700 | 1 340 000 | 132 400 | 312 000 |
| KGR-15 | Val-Glu-Ala-Asp | 337 000 | 34 400 | 306 100 | 338 000 | 47 800 | 2 222 000 | 130 600 | 308 000 |
| KGR-17 | Nle-Glu-Thr-Asp | 487 000 | 52 600 | 12 000 | 72 000 | 8 900 | 664 000 | 62 100 | 164 000 |
| KGR-18 | Leu-Glu-Val-Asp | 36 400 | 6 400 | 457 000 | 652 000 | 97 000 | 1 950 000 | 69 200 | 377 000 |
| KGR-19 | Leu-Gln-Ser-Asp | 359 000 | 36 400 | 28 800 | 21 300 | 2 000 | 1 114 000 | 80 300 | 315 000 |
| KGR-20 | Thr-Glu-Ser-Asp | 5 200 | 1 800 | 92 100 | 195 000 | 24 500 | 658 000 | 9 600 | 66 000 |
| KGR-23 | Phe(2-Cl)-Glu-Ala-Asp | 3 725 000 | 60 200 | 255 000 | 58 400 | 58 400 | 2 340 000 | 104 200 | 374 000 |
| KGR-25 | Phe(4-NH<sub>2</sub>)-hGlu-Cha-Asp | 90 300 | 2 400 | 80 500 | 89 900 | 16 600 | 374 000 | 9 200 | 97 600 |
| KGR-26 | Leu-Phe(4-NH<sub>2</sub>)-Aoc-Asp | 51 800 | 20 800 | 146 000 | 59 200 | 22 600 | 918 000 | 76 400 | 144 000 |
| KGR-28 | Leu-Abu-hCha-Asp | 199 500 | 12 800 | 104 000 | 123 000 | 11 200 | 2 352 000 | 183 000 | 148 000 |
| KGR-29 | Oic-Glu-hPro-Asp | 16 800 | 22 200 | 7 300 | 1 100 | 500 | 882 000 | 73 100 | 89 800 |
| KGR-30 | Val-Glu-Lys(Me)-Asp | 12 600 | 9 600 | 114 600 | 66 900 | 10 500 | 530 000 | 8 700 | 63 900 |
| KGR-31 | Val-hSer-Val-Asp | 800 | 100 | 41 800 | 19 700 | 15 300 | 498 000 | 7 900 | 10 800 |
| KGR-32 | DhPhe-Glu-Gln-Asp | 20 300 | 440 | 12 000 | 1 300 | 2 800 | 134 000 | 11 500 | 22 700 |
| KGR-33 | DChg-hGlu-hCha-Asp | 71 500 | 1 200 | 26 900 | 8 000 | 6 200 | 966 000 | 82 500 | 154 000 |
| KGR-34 | Leu-Glu-Thr(Bzl)-Asp | 63 000 | 21 400 | 296 500 | 446 000 | 75 200 | 832 000 | 42 300 | 440 000 |
| KGR-35 | Ser-Tle-Pro-Asp | 155 000 | 18 400 | 229 400 | 1 300 | 161 600 | 1 756 000 | 75 700 | 137 000 |
| KGR-36 | Oic-Nle-Pro-Asp | 20 300 | 1 400 | 6 300 | 600 | 700 | 104 000 | 66 800 | 12 000 |
| KGR-37 | hPro-Leu-Ala-Asp | 424 000 | 11 200 | 13 300 | 600 | 7 000 | 2 900 000 | 62 200 | 177 000 |
| KGR-38 | Lys(Ac)-hGlu-Ala-Asp | 265 000 | 20 400 | 27 800 | 3 550 | 3 000 | 990 000 | 84 800 | 24 500 |
| KGR-39 | Pro-Tle-Gly-Asp | 9 800 | 400 | 16 700 | 500 | 3 800 | 310 000 | 59 900 | 15 400 |
| KGR-40 | Oic-Leu-Ala-Asp | 61 000 | 1 300 | 10 400 | 500 | 1 800 | 422 000 | 113 700 | 8 200 |
| KGR-41 | Ser-Phe-Ala-Asp | 25 200 | 800 | 263 000 | 1 100 | 48 700 | 502 000 | 55 200 | 115 000 |
| KGR-42 | Nle-hGlu-Pro-Asp | 783 000 | 51 400 | 98 000 | 5 700 | 12 300 | 2 284 000 | 106 200 | 322 000 |
| KGR-43 | Leu-Tle-Tyr-Asp | 9 100 | 300 | 33 900 | 24 100 | 600 | 277 000 | 27 900 | 407 000 |
| KGR-44 | Pro-Gln-Ala-Asp | 249 000 | 7 600 | 37 500 | 4 800 | 6 900 | 145 000 | 120 200 | 143 000 |
| KGR-45 | Asp-Ala-Phe(NH<sub>2</sub>)-Asp | 26 700 | 1 800 | 1 910 000 | 113 000 | 216 000 | 57 000 | 4 800 | 46 000 |
| KGR-46 | Asp-Val-Thr(Bzl)-Asp | 58 000 | 1 700 | 5 213 000 | 30 800 | 1 010 000 | 1 164 000 | 8 700 | 393 000 |
| KGR-47 | Asp-hGlu-Leu-Asp | 45 000 | 23 400 | 4 145 000 | 243 000 | 610 800 | 924 000 | 9 100 | 260 000 |
| KGR-49 | Trp-Lys-Glu-Asp | 684 000 | 20 400 | 6 500 | 1 900 | 18 700 | 43 000 | 2 100 | 14 500 |
| KGR-50 | Tyr-Lys-Ala-Asp | 1 240 000 | 14 200 | 8 600 | 200 | 2 200 | 32 000 | 10 800 | 123 000 |
| KGR-51 | Tyr-Thz-Ala-Asp | 88 000 | 900 | 9 500 | 300 | 1 700 | 28 000 | 2 200 | 6 600 |
| KGR-52 | Trp-Arg-Ser-Asp | 2 340 000 | 16 600 | 6 200 | 200 | 300 | 31 000 | 3 900 | 28 400 |
| KGR-53 | Tle-hGlu-hCha-Asp | 127 000 | 30 000 | 89 500 | 279 000 | 7 000 | 1 426 000 | 86 200 | 484 000 |
| KGR-54 | Ser-Tle-Abu-Asp | 32 200 | 2 200 | 397 000 | 12 800 | 107 800 | 790 000 | 57 000 | 150 000 |
| KGR-55 | Phe(4-I)-Glu-Thr-Asp | 1 850 000 | 50 000 | 156 000 | 44 100 | 51 400 | 796 000 | 28 200 | 154 000 |
| KGR-56 | Phe(4-Me)-hGlu-Tle-Asp | 49 000 | 1 700 | 132 000 | 5 300 | 9 000 | 167 000 | 3 500 | 218 000 |
| KGR-57 | Phe(4-I)-Glu-Ile-Asp | 252 000 | 12 800 | 197 000 | 37 300 | 64 300 | 956 000 | 58 100 | 235 000 |
| KGR-58 | Val-Lys-Gln-Asp | 3 000 | 300 | 9 100 | 2 000 | 800 | 162 000 | 14 600 | 55 000 |
| KGR-59 | Tle-Gln-Ser-Asp | 16 000 | 500 | 36 900 | 25 100 | 2 000 | 1 426 000 | 54 700 | 314 000 |
| KGR-60 | DhPhe-hGlu-hPro-Asp | 954 000 | 24 000 | 6 000 | 400 | 300 | 890 000 | 53 300 | 12 400 |
| KGR-61 | Chg-Cit-hSer-Asp | 5 600 | 200 | 7 600 | 500 | 900 | 48 000 | 16 200 | 25 800 |
| KGR-63 | hPro-Val-Ser-Asp | 36 400 | 600 | 11 700 | 600 | 2 500 | 431 000 | 31 400 | 63 000 |
| KGR-64 | hLeu-Glu-Ala-Asp | 1 140 000 | 66 800 | 75 500 | 13 500 | 9 600 | 978 000 | 166 000 | 261 000 |
| KGR-65 | Val-Tle-Abu-Asp | 328 000 | 11 900 | 181 900 | 39 300 | 25 300 | 1 506 000 | 156 700 | 259 000 |
| KGR-66 | hPro-Tle-Ser-Asp | 67 000 | 4 500 | 9 600 | 900 | 4 100 | 1 258 000 | 65 900 | 260 000 |
| KGR-67 | Leu-Phe-Ser-Asp | 34 000 | 3 300 | 17 700 | 3 000 | 900 | 82 000 | 55 000 | 355 000 |
| KGR-68 | Cha-Glu-Aoc-Asp | 254 000 | 19 800 | 75 000 | 21 100 | 11 100 | 406 000 | 68 600 | 92 400 |
| KGR-69 | Nle-Glu-Ser-Asp | 1 015 000 | 60 200 | 29 200 | 48 000 | 3 500 | 1 038 000 | 78 000 | 381 000 |
| KGR-70 | hLeu-Tle-Ser-Asp | 987 000 | 31 600 | 6 600 | 400 | 600 | 42 000 | 48 100 | 178 000 |
| KGR-72 | Trp-Glu-Ala-Asp | 4 870 000 | 74 800 | 46 700 | 18 500 | 20 400 | 1 890 000 | 103 500 | 301 000 |
| KGR-73 | hleu-hGlu-Ser-Asp | 897 000 | 31 600 | 27 400 | 3 800 | 3 800 | 24 000 | 51 500 | 292 000 |
| KGR-74 | Asp-Thr-Abu-Asp | 32 000 | 16 600 | 2 041 000 | 31 500 | 363 200 | 141 000 | 22 600 | 122 000 |
| KGR-75 | Asp-Glu-Leu-Asp | 36 000 | 32 800 | 3 727 000 | 327 000 | 574 000 | 798 000 | 22 400 | 126 000 |
| KGR-77 | Asp-Ser-Ala-Asp | 7 600 | 9 600 | 865 000 | 8 100 | 156 000 | 84 000 | 4 300 | 96 000 |
| KGR-78 | Asp-2Nal-Ala-Asp | 23 100 | 1 200 | 1 367 000 | 1 600 | 197 000 | 108 000 | 24 300 | 118 000 |
| KGR-79 | Asp-Nle-Thr(Bzl)-Asp | 45 000 | 2 600 | 4 622 000 | 27 000 | 938 000 | 1 434 000 | 4 700 | 235 000 |
| KGR-80 | Asp-hGlu-Ile-Asp | 36 400 | 22 600 | 3 106 000 | 160 000 | 488 000 | 730 000 | 6 100 | 101 000 |
| KGR-81 | Asp-Leu-Abu-Asp | 46 200 | 16 600 | 1 792 000 | 11 500 | 367 600 | 1 152 000 | 22 200 | 137 000 |
| KGR-82 | Asp-Ser-Val-Asp | 14 700 | 17 200 | 2 624 000 | 85 000 | 517 200 | 82 000 | 1 200 | 58 400 |
| KGR-83 | Asp-Phe(4-I)-Ala-Asp | 30 100 | 14 800 | 1 440 000 | 3 200 | 190 400 | 34 000 | 22 100 | 246 000 |
| KGR-85 | Asp-Dap(Z)-Ile-Asp | 30 800 | 21 200 | 5 484 000 | 251 000 | 928 000 | 548 000 | 5 100 | 195 000 |
| KGR-87 | Asp-Phe-Tle-Asp | 3 600 | 200 | 179 000 | 3 800 | 27 500 | 121 000 | 8 900 | 32 000 |
| KGR-88 | Asp-Leu-Ala-Asp | 35 400 | 2 300 | 472 000 | 1 400 | 169 800 | 246 000 | 21 800 | 187 000 |
| KGR-89 | Asp-hGlu-Oic-Asp | 71 000 | 23 500 | 1 789 000 | 80 000 | 320 200 | 832 000 | 31 300 | 263 000 |
| KGR-90 | Leu-Dab-Ser(Bzl)-Asp | 21 700 | 1 800 | 2 742 000 | 13 300 | 393 000 | 442 000 | 2 100 | 120 000 |
| KGR-91 | Asp-hGlu-Oic-Asp | 78 400 | 46 400 | 2 114 000 | 57 700 | 355 400 | 188 000 | 35 600 | 231 000 |
| KGR-92 | Asp-Nva-Oic-Asp | 22 400 | 17 600 | 4 793 000 | 23 800 | 940 000 | 524 000 | 25 700 | 165 000 |
| KGR-93 | Asp-Thr(Bzl)-Ala-Asp | 39 200 | 1 800 | 2 066 000 | 25 900 | 557 400 | 191 000 | 29 600 | 304 000 |
| KGR-94 | Asp-Nle-Oic-Asp | 30 100 | 1 500 | 1 669 000 | 16 000 | 162 000 | 90 000 | 16 500 | 118 000 |
| KGR-96 | Gln-Leu-Pro-Asp | 19 000 | 900 | 58 900 | 600 | 24 800 | 44 000 | 21 800 | 14 000 |
| KGR-97 | Leu-hGlu-hSer-Asp | 8 400 | 400 | 187 000 | 16 200 | 16 100 | 97 000 | 3 600 | 36 600 |
| KGR-98 | Val-hGlu-Glu(Chx)-Asp | 34 000 | 3 400 | 292 000 | 29 500 | 23 300 | 139 000 | 42 500 | 85 000 |
| KGR-99 | Ser-Tle-Glu-Val-Asp | 26 900 | 34 800 | 714 000 | 421 000 | 47 000 | 758 000 | 55 100 | 207 000 |
| KGR-100 | Asp-Leu-Thr-Ala-Asp | 72 000 | 5 400 | 88 500 | 5 100 | 3 100 | 210 000 | 70 300 | 144 000 |
| KGR-103 | Phe-Asp-Glu-Pro-Asp | 68 500 | 46 000 | 3 297 000 | 86 600 | 644 000 | 1 514 000 | 45 400 | 177 000 |
| KGR-104 | Leu-Gln-Glu-Asp | 26 600 | 38 600 | 17 700 | 9 600 | 400 | 364 000 | 39 300 | 242 000 |
| KGR-105 | Leu-Dab-Ser(Bzl)-Asp | 21 000 | 1 300 | 15 000 | 4 900 | 1 600 | 192 000 | 8 800 | 103 000 |
| KGR-106 | Nle-hGlu-hPhe-Asp | 675 000 | 59 400 | 110 000 | 683 000 | 11 800 | 1 564 000 | 103 600 | 399 000 |
| KGR-107 | Gln-Leu-hGlu-Pro-Asp | 834 000 | 71 000 | 421 000 | 24 600 | 67 000 | 1 134 000 | 117 300 | 194 000 |

**Extended Data Table 3.**
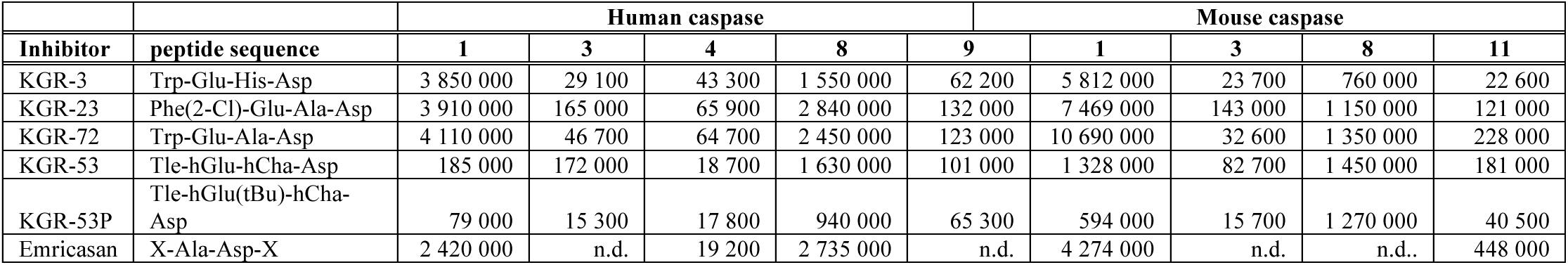
Caspase k_obs_/I values (M^-1^s^-1^) for caspase inhibitors having the format Ac-P4-P3-P2-P1(Asp)-AOMK, or emricasan (N-[2-(1,1-dimethylethyl)phenyl]-2-oxoglycyl-N-[(1S)-1-(carboxymethyl)-2-oxo-3-(2,3,5,6-tetrafluorophenoxy)propyl]-L-alaninamide). See methods for how this parameter was calculated. n.d., not determined.

**Extended Data Table 4.** Apparent permeability (Papp) values for a subset of tetrapeptide caspase inhibitors in the apical to basolateral direction (A-to-B) using gMDCKI cells. Media in the apical and basolateral chambers were pH 7.4. Results are the mean of at least 4 technical replicates.

| Inhibitor | Concentration (uM) | Incubation Time (min) | A-to-B Papp (x10<sup>-06</sup> cm/s) | Intracellular Concentration (uM) | Mean Mass Balance (%) without Cell Lysate | Permeability Classification |
| --- | --- | --- | --- | --- | --- | --- |
| KGR-3 | 25 | 390 | 0.0215 | 0.614 | 89.5 | Very low |
| KGR-53 | 25 | 390 | 0.0249 | Below limit of detection | 86.5 | Very low |
| KGR-53P | 25 | 390 | 0.0208 | Below limit of detection | 86.5 | Very low |
| KGR-72 | 25 | 390 | 0.0200 | Below limit of detection | 91.1 | Very low |

