## Supplemental Figures for "Delivery of caspase inhibitors through GSDMD pores to inhibit pyroptosis"

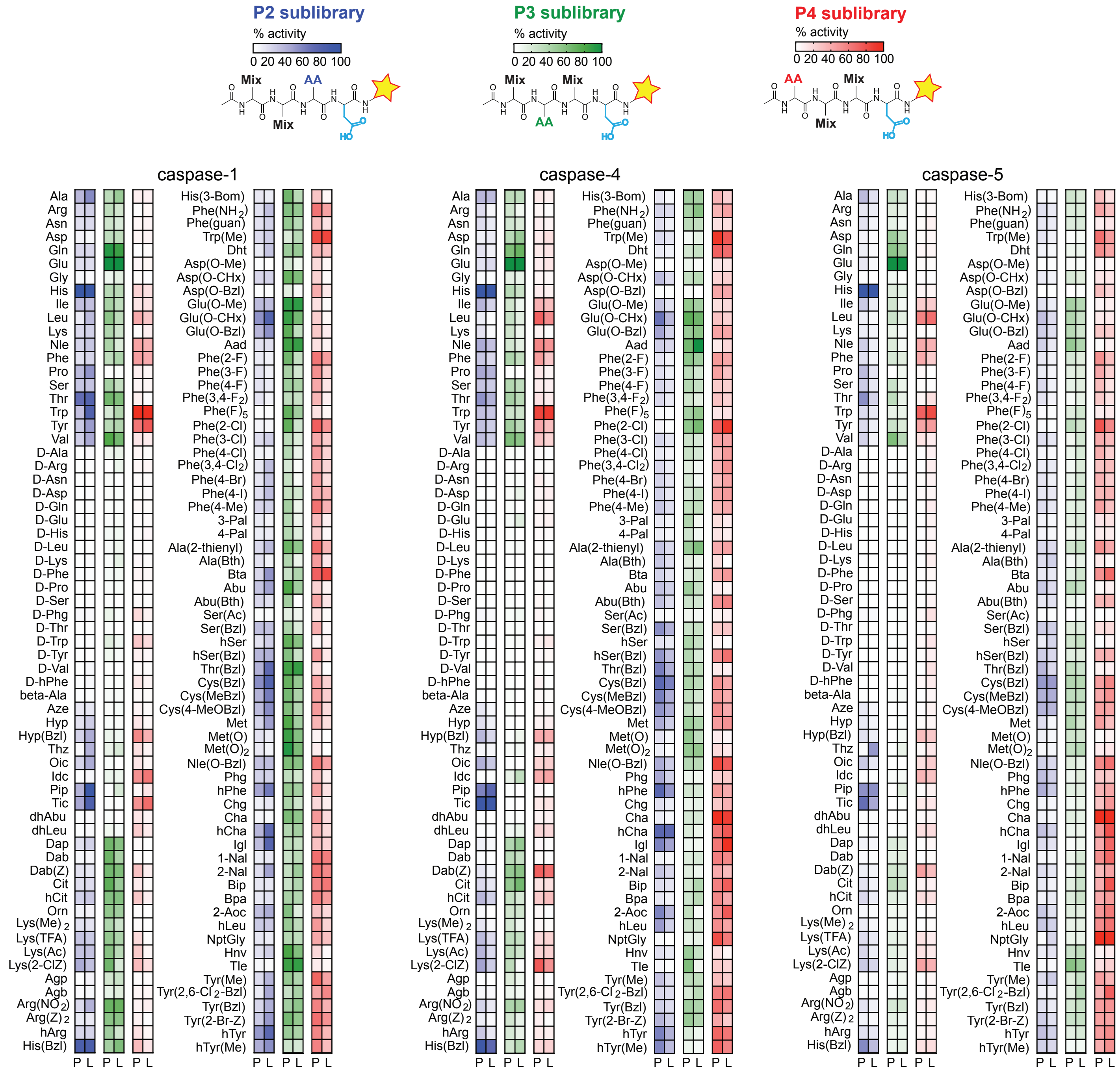


**Extended Data Fig. 1. Substrate specificity screening results of human caspase-1, -4, and -5**.
Hybrid Combinatorial Substrate Libraries were screened using purified recombinant enzyme (P) or *E.coli* lysates (L) containing the caspases.
Mix: equimolar mixture of 19 natural amino acids, AA: fixed natural or unnatural amino acid ☆: reporter group (ACC).

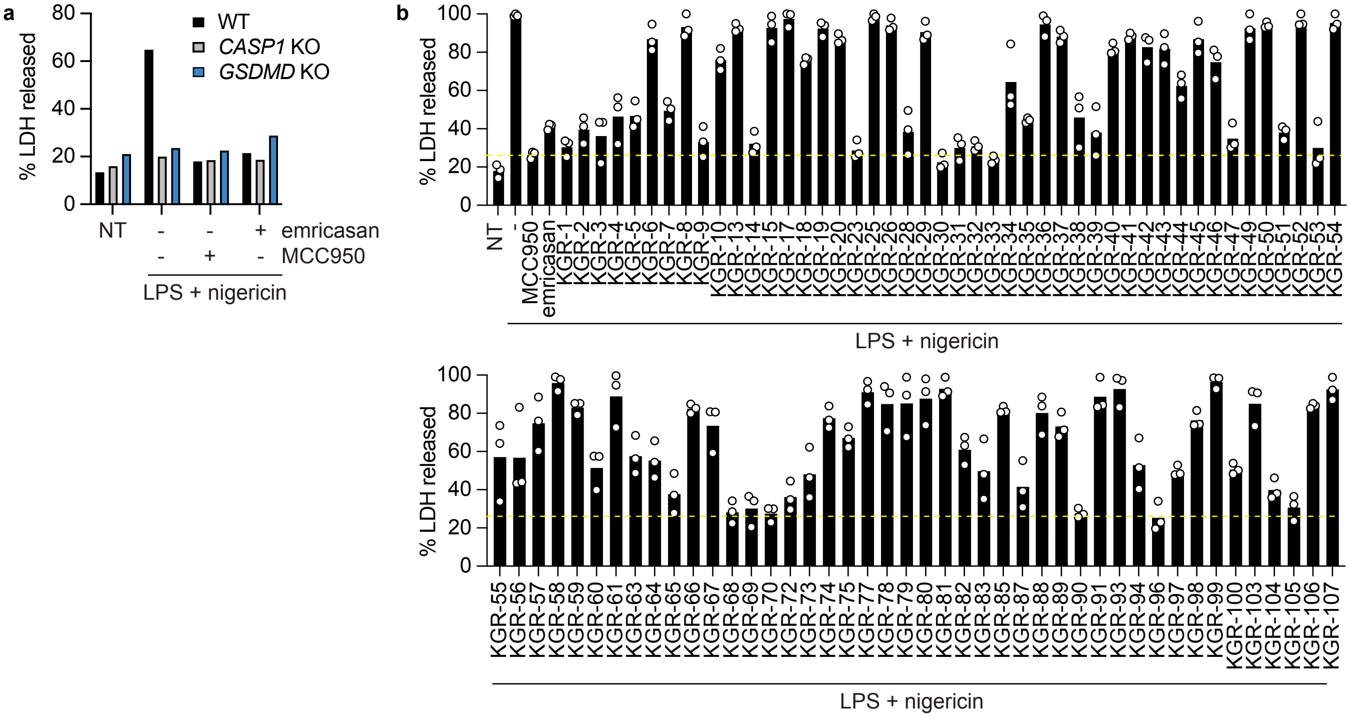


**Extended Data Fig. 2. Screening for inhibitors of nigericin-induced pyroptosis in THP-1 cells.**

**a and b,** Percentage of LDH released from LPS-primed THP-1 cells after treatment with 6.25 (a) or 5 (b) μM nigericin for 2 h. Where indicated, cells were co-treated with 25 μM MCC950, 25 μM emricasan, or 25 μM tetrapeptide-AOMK (assigned the prefix KGR for tracking purposes). NT, no treatment. WT, wild-type. KO, knockout. The horizontal, dashed yellow line in (b) indicates the degree of protection provided by MCC950. Results in (a) are representative of 3 independent experiments. Bars in (b) indicate the mean of 3 independent experiments represented by circles.

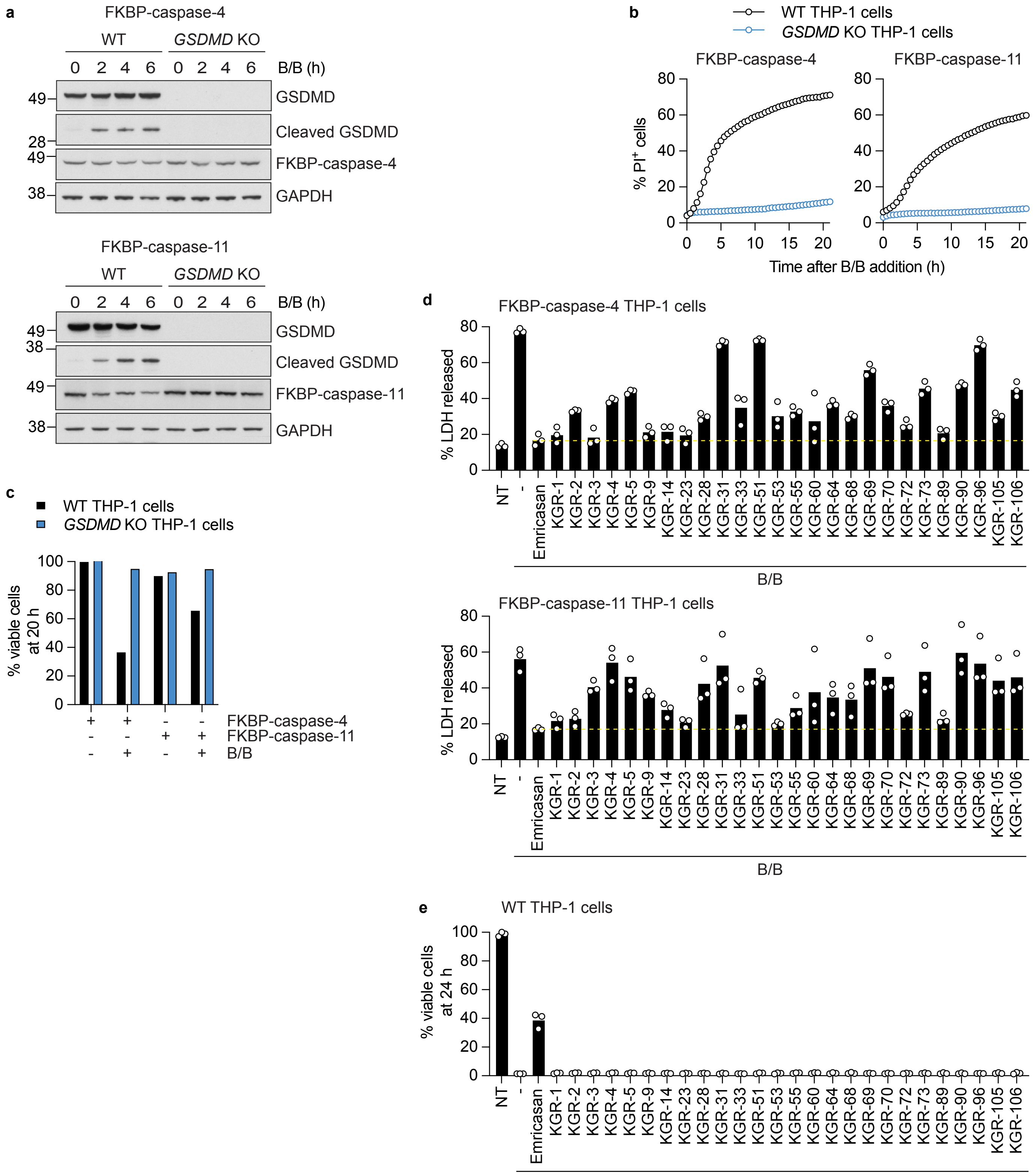


**Extended Data Fig. 3. Screening for inhibitors of caspase-4 or -11.**

**a,** Western blots of THP-1 cells treated with 50 nm B/B. Results representative of 2 independent experiments. **b,** Percentage of THP-1 cells expressing FKBP-caspase-4 (left) or FKPP-caspase-11 (right) that exhibited nuclear PI staining after treatment with 50 nM B/B. Results representative of 3 independent experiments. **c,** Percentage of viable THP-1 cells as measured by CellTiter-Glo assay after treatment with 50 nM B/B for 20 h. Results representative of 3 independent experiments. **d,** Percentage of LDH released from THP-1 cells after treatment with 50 nM B/B for 24 h. Where indicated, cells were co-treated with 10 μM emricasan or tetrapeptide-AOMK. NT, no treatment. **e,** Percentage of viable THP-1 cells after 24 h as measured by CellTiter-Glo assay. Where indicated, cells were treated with 1 μM staurosporine ± 10 μM tetrapeptide-AOMK. Bars in (d) and (e) indicate the mean of 3 independent experiments represented by circles. The horizontal, dashed yellow lines indicate the degree of protection provided by emricasan.

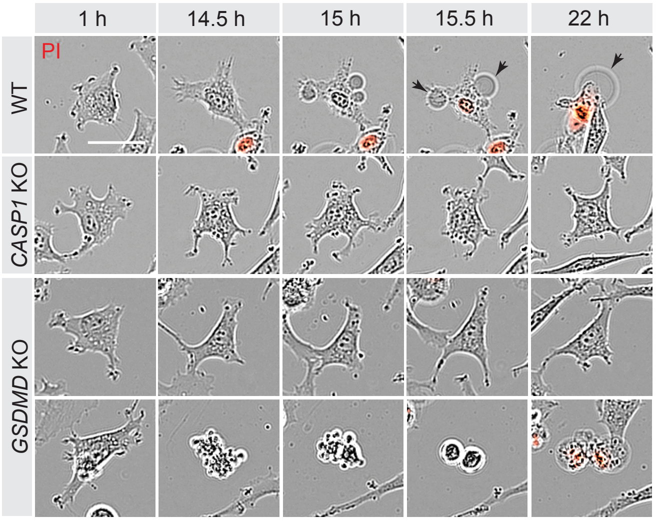


**Extended Data Fig. 4. VBP induces apoptosis and secondary membrane rupture in *GSDMD* KO Ea.hy926 cells.**
Ea.hy926 cells treated with 20 μM VBP in the presence of 1 μg/ml PI. Scale bar, 40 μm. In cultures of *GSDMD* KO cells, most VBP-treated cells remained viable out to 22 h (top row), but some cells exhibited apoptotic blebbing with subsequent membrane rupture and PI uptake (bottom row). Scale bar, 40 μm. Black arrows indicate ballooning pyroptotic cells. Results representative of 3 independent experiments.

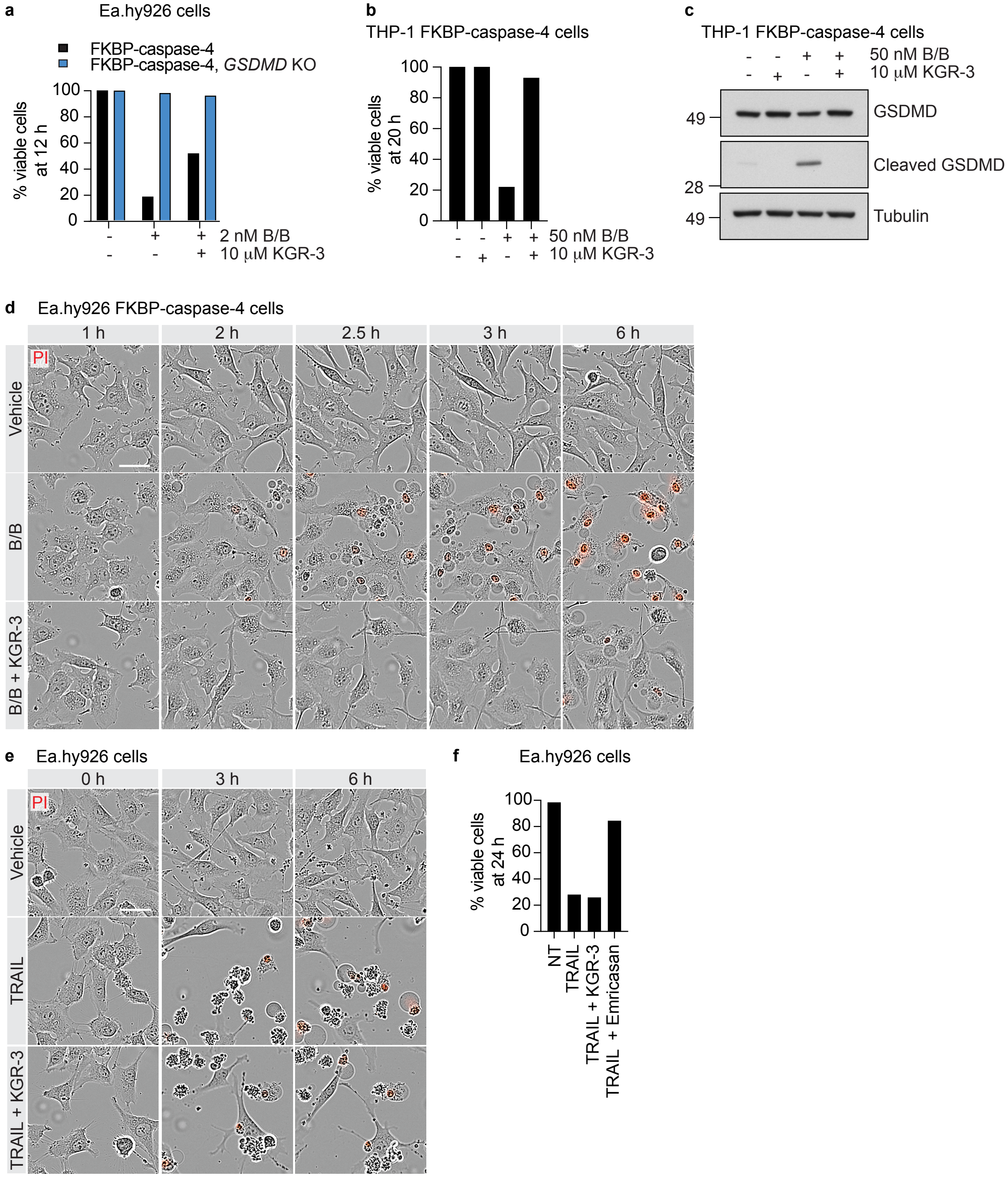


**Extended Data Fig. 5. KGR-3 suppresses caspase-4-driven pyroptosis, but not caspase-8-driven apoptosis.**
**a and b,** Percentage of viable cells as measured by CellTiter-Glo assay. Results representative of 5 (a) and 2 (b) independent experiments. **c,** Western blots of THP-1 cells expressing FKBP-caspase-4 and treated as indicated for 6 h. Results representative of 2 independent experiments. **d,** Ea.hy926 cells expressing FKBP-caspase-4 and treated with 2 nM B/B ± 10 μM KGR-3 in the presence of 1 μg/ml PI. Scale bar, 40 μm. PI+ cells are red. Results representative of 3 independent experiments. **e,** Ea.hy926 cells treated with 100 ng/ml TRAIL ± 10 μM KGR-3 in the presence of 1 μg/ml PI. Scale bar, 40 μm. PI+ cells are red. Results representative of 3 independent experiments. **f,** Percentage of viable Ea.hy926 cells after treatment with 100 ng/ml TRAIL ± 10 μM KGR-3 or emricasan. Results representative of 3 independent experiments. NT, no treatment.

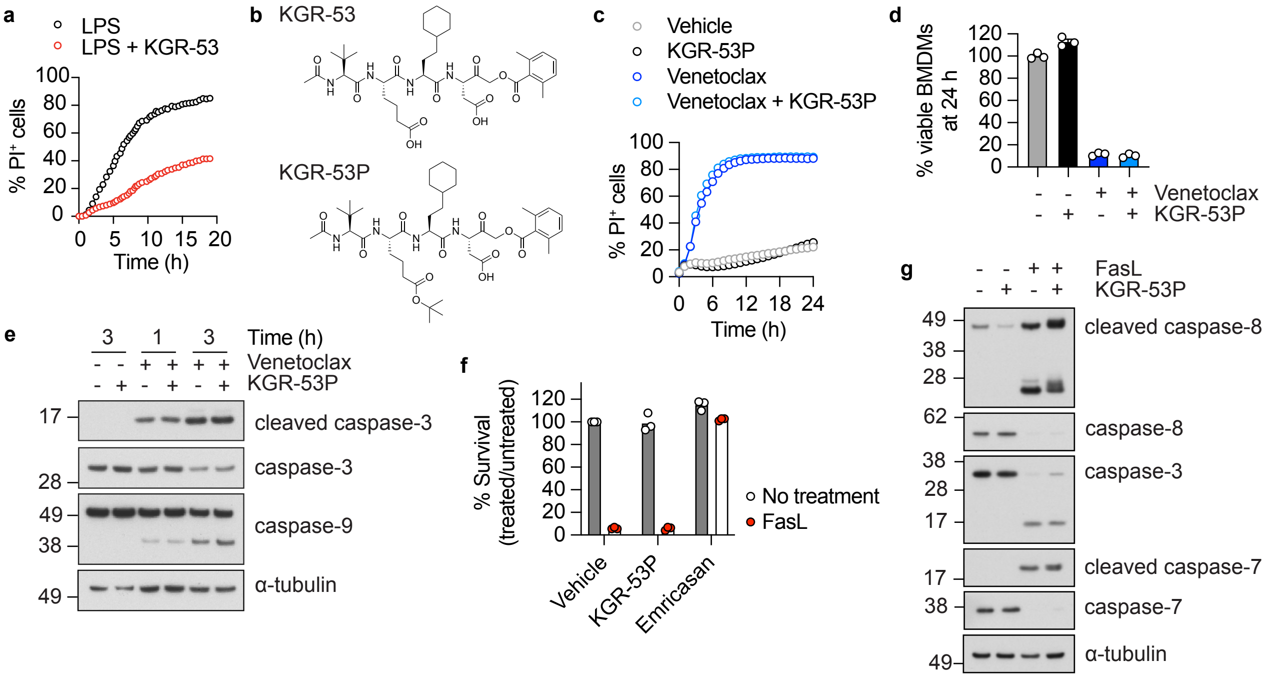


**Extended Data Fig. 6. Characterization of KGR-53 and KGR-53P.**

**a,** Percentage of LPS-transfected BMDMs that exhibited nuclear PI staining after treatment with 30 µM KGR-53. **b,** Structures of KGR-53 and KGR-53P. **c,** Percentage of BMDMs with nuclear PI staining after treatment with 25 µM Venetoclax ± 10 µM KGR-53P. **d,** Percentage of viable BMDMs by CellTiter-Glo assay after treatment with 25 µM Venetoclax ± 10 µM KGR-53P. **e**, Western blots of BMDMs treated with 25 µM Venetoclax ± 10 µM KGR-53P. Results representative of 2 independent experiments. **f**, Percentage of PI-negative thymocytes after treatment with 100 ng/ml FasL ± 10 µM KGR-53P or emricasan for 20 h. Circles, cells from different mice (n = 3). **g**, Western blots of thymocytes treated with 100 ng/ml FasL ± 10 µM KGR-53P for 2 h.
